# Development of a new class of liver receptor homolog-1 (LRH-1) agonists by photoredox conjugate addition

**DOI:** 10.1101/2020.03.16.994400

**Authors:** Jeffery L. Cornelison, Michael L. Cato, Alyssa M. Johnson, Emma H. D’Agostino, Diana Melchers, Anamika B. Patel, Suzanne G. Mays, René Houtman, Eric A. Ortlund, Nathan T. Jui

## Abstract

LRH-1 is a nuclear receptor that regulates lipid metabolism and homeostasis, making it an attractive target for the treatment of diabetes and non-alcoholic fatty liver disease. Building on recent structural information about ligand binding from our labs, we have designed a series of new LRH-1 agonists that further engage LRH-1 through added polar interactions. While the current synthetic approach to this scaffold has, in large part, allowed for decoration of the agonist core, significant variation of the bridgehead substituent is mechanistically precluded. We have developed a new synthetic approach to overcome this limitation, identified that bridgehead substitution is necessary for LRH-1 activation, and described an alternative class of bridgehead substituents for effective LRH-1 agonist development. We determined the crystal structure of LRH-1 bound to a bridgehead-modified compound, revealing a promising opportunity to target novel regions of the ligand-binding pocket to alter LRH-1 target gene expression.

---

Liver receptor homolog-1 (LRH-1) is a member of the nuclear receptor (NR) family of ligand-regulated transcription factors that sense lipophilic signaling molecules and produce biological responses. LRH-1 regulates a variety of cellular and organismal processes, including reverse cholesterol transport,^1^ steroidogenesis,^2^ endoplasmic reticulum stress resolution,^3^ intestinal cell renewal,^4^ embryonic development,^5^ and bile acid biosynthesis.^6–8^ Its roles in lipid and glucose homeostasis have drawn attention to LRH-1 as a potential target for treating type II diabetes and non-alcoholic fatty liver disease (NAFLD),^6^ while its role in intestinal cell renewal has made it a promising target for the treatment of inflammatory bowel disease.^9^ Therefore, compounds that modulate LRH-1 activity could be valuable for the treatment of multiple diseases.

Dietary phospholipids are the putative endogenous ligands for LRH-1,^10^ and a number of studies demonstrated that phosphatidylcholines such as diundecanoylphosphatidylcholine (DUPC) and dilauroylphosphatidylcholine (DLPC) preferentially activate LRH-1.^6,11,12^ However, because of the low potency and poor physicochemical properties of phospholipids, effective synthetic probes are required for characterization of LRH-1 biology. Towards this end, several laboratories have madesignificant advances in developing potent LRH-1 modulators.^13–15^ Despite these advances, rational design has been difficult, in partbecause of the large, highly hydrophobic LRH-1 ligand binding pocket. Due to this lipophilicity and a scarcity of sites for anchoring polar interactions, even highly similar compounds can bind unpredictably,^15,16^ further complicating systematic agonist development.

Recently, our lab has identified key anchoring interactions in the binding pocket that established the mechanism of binding for the privileged [3.3.0] bicyclic hexahydropentalene (6HP) substructure (shown in Fig. 1, top), which was first identified by Whitby.^17,18^ Employing this information has led to the design of more potent LRH-1 agonists through optimization of R^1^ (**“6N”** | EC_50_ = 15 ± 8 nM; Fig. 2, left),^19^ as well as more strongly activating agonists, through optimization of R^2^ (**“6HP-CA”** | 2.3 ± 0.2-fold activation over vehicle; Fig. 2, right).^20^ To further interrogate the structural requirements for LRH-1 activation by 6HP agonists, we sought to vary the bridgehead substituent (R^3^) (Fig. 1, bottom). Because Whitby’s 3-component cyclization results in either heteroatom- or vinyl-substitution at this position,^21,22^ we considered the alternative synthetic approach that is outlined in Fig. 1, where the installation of R^3^ would be accomplished through functionalization of tetrasubstituted olefin **1**. Although alkenes of this type are notoriously unreactive,^23^ this plan was appealing because it would allow for modular variation (or deletion) of R^3^, and regioselective enol-triflate formation would allow for installation of different alkyl tails (R^2^).

**Fig. 1.**
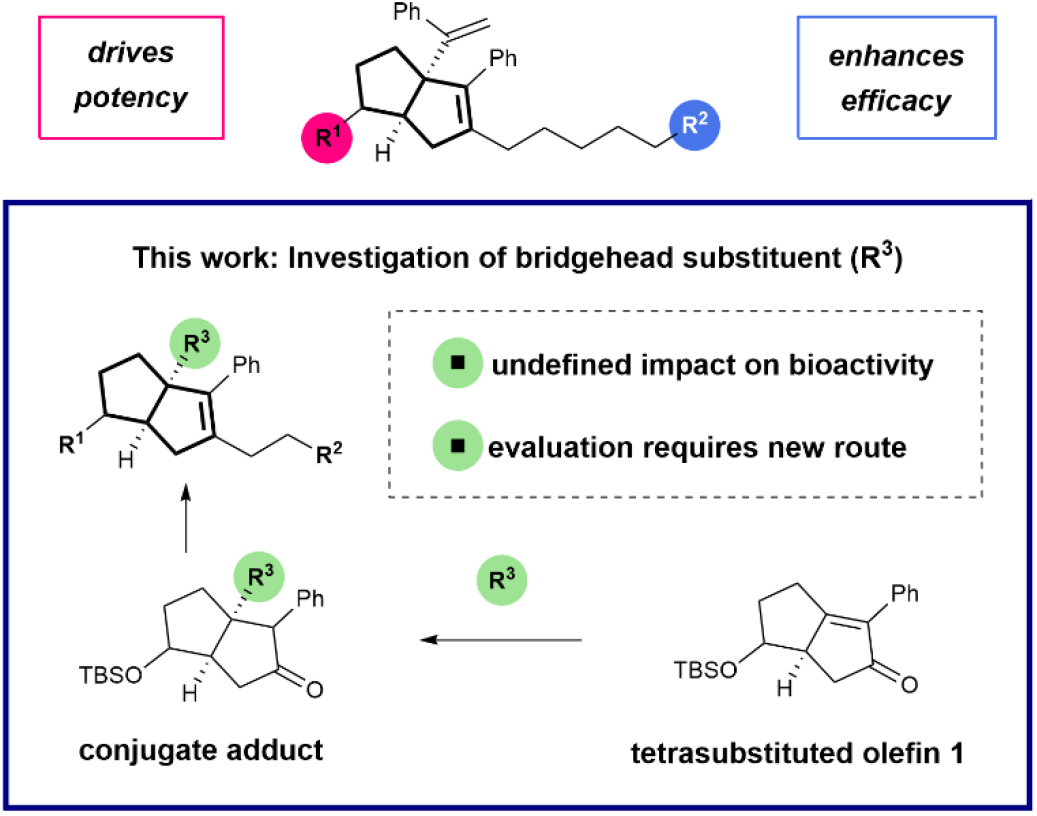
Design of Hexahydropentalene LRH-1 agonists

**Fig. 2.**
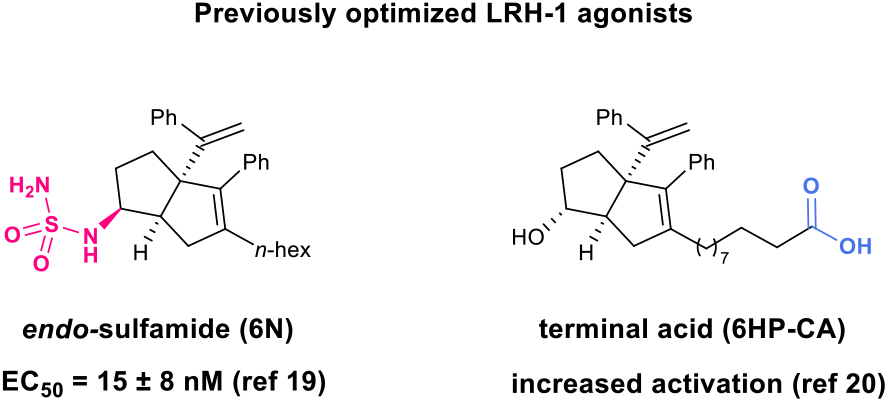
LRH-1 agonists previously reported by our lab with key polar groups highlighted

To evaluate the feasibility of this proposal, we prepared olefin **1** through Pauson-Khand cyclization of an appropriately-substituted 1,6-enyne (see SI for details). Because the parent styryl R^3^ group does not appear to make any critical contacts in the ligand binding site of LRH-1, we first sought to investigate the necessity of substitution at this position. Accordingly, our synthetic plan involved reduction of the enone function, followed by elaboration of the resulting material to the corresponding *endo*-sulfamide or terminal acid analogs, such that direct comparison with either of the parent compounds would reveal the importance of R^3^. While a range of reducing conditions were able to engage **1**, we found that the cleanest profile was observed in the presence of palladium on carbon and sodium borohydride, followed by *in situ* triflation to afford **2** as a single regioisomer (69% yield, over two steps). This product could be utilized in a Negishi coupling under Knochel conditions,^24^ where the SPhos-supported palladium catalyst afforded methyl decanoate derivative **3**. Routine silyl ether cleavage and saponification gave rise to **5**, the direct analog of 6HP-CA lacking the bridgehead styrene.

## Scheme 1.

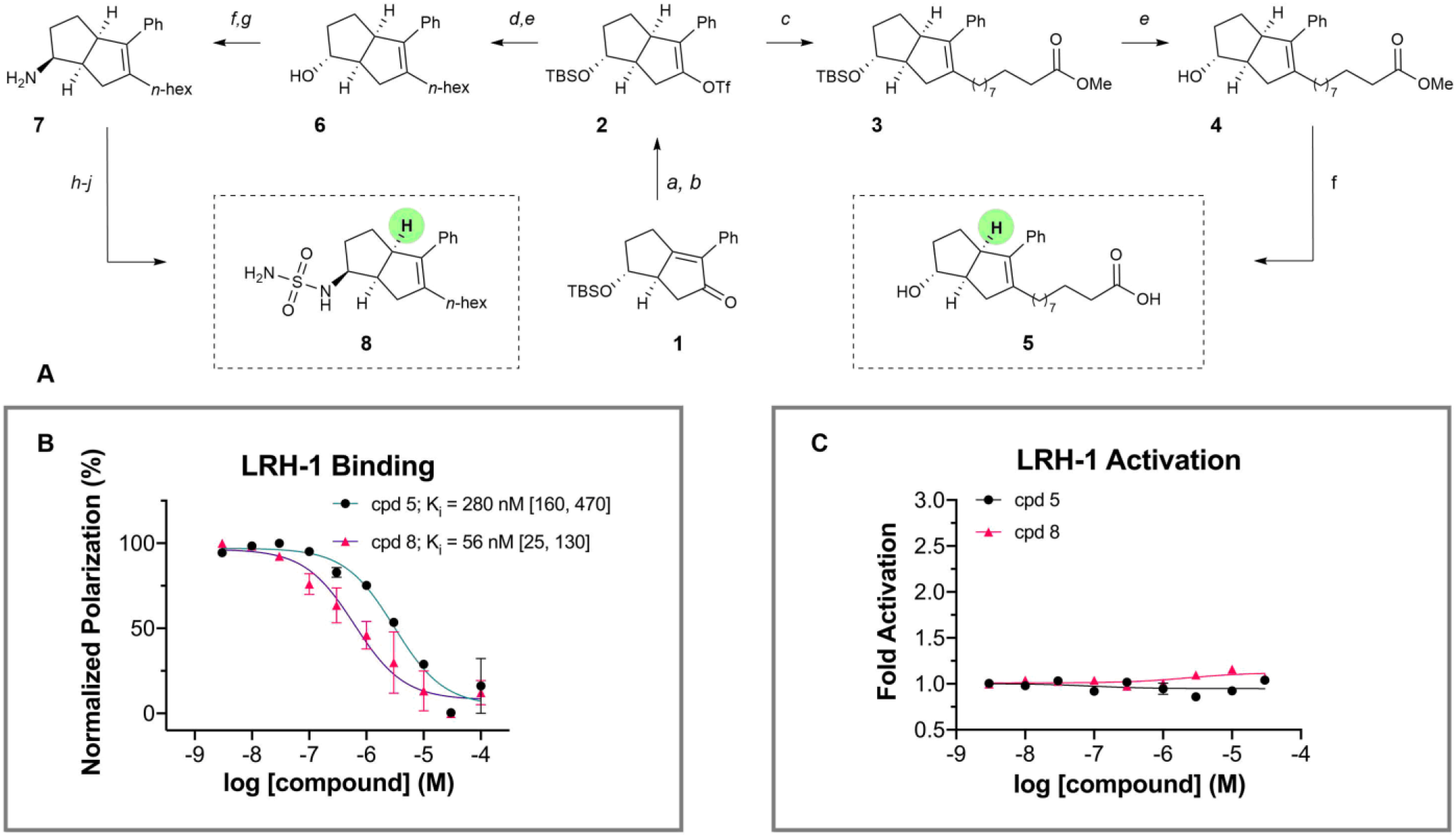
Synthesis and Evaluation of Unsubstituted Bridgehead Compounds **A**) Reagents and conditions: (a) Pd/C, NaBH_4_, AcOH, PhMe, 23 °C; 1h; (b) NaH, PhNTf_2_, 0 – 23 °C; 69% yield over two steps; (c) SPhos Pd G3, SPhos, IZn(CH_2_)_9_CO_2_Me·LiCl, THF, 50 °C; 16 h; (d) SPhos Pd G3, SPhos, IZn(CH_2_)_5_CH_3_·LiCl, THF, 50 °C; 16 h; (e) conc. aq. HCl, MeOH, 23 °C; 1 h, 41 – 43% yield over two steps; (f) LiOH, H_2_O, THF, 50 °C; 16 h, 97% yield; (g) TPAP, NMO, MeCN, 23°C; 1 h, 81% yield; (h) i. Ti(OiPr)_4_, NH_3_, MeOH, 23 °C; 5 h; ii. NaBH_4_, MeOH, 23 °C; 5 h, 59% yield; (i)chlorosulfonylisocyanate, *^t^*BuOH, TEA, DCM, 0 – 23 °C; 1.5 h, 31% yield; (j) conc. aq. HCl, dioxane, 0 – 40 °C; 14 h, 77% yield. **B**) Fluorescence polarization (FP) evaluation of compounds **5** and **8**. Data shown as mean ± SEM from two independent experiments. K_i_ values are given with 95% confidence intervals in brackets. **C**) Luciferase reporter data for **5** and **8** shown as mean ± SEM from three biological replicates.

To access the simplified *endo*-sulfamide analog, vinyl triflate **2** was reacted with hexylzinc iodide under the same coupling conditions, which after acidic alcohol deprotection, afforded **6** in moderate yield (43%). Ley oxidation gave the corresponding ketone, which underwent highly diastereoselective reductive amination with ammonia to afford the *endo* amine **7**. Sulfamide installation was accomplished using chlorosulfonylisocyanate and *tert*-butanol, followed by acidic decomposition of the resulting N-Boc sulfamide to afford analog **8**.

We tested the biological activity of this simplified 6HP series using a fluorescence polarization (FP) competition ligand binding assay recently developed in our lab^25^ and a luciferase reporter assay to measure LRH-1 transcriptional activity. We assessed the binding affinity (K_i_), in-cell potency (EC_50_), and efficacy (fold activation) of compounds **5** and **8**, the direct analogs of **6HP-CA** and **6N** respectively, with the bridgehead group entirely removed. The compound containing a sulfamide anchoring group (**8**) demonstrated low nanomolar binding affinity (K_i_ = 56 nM), while the compound with a charged tail (**5**) demonstrated mid nanomolar affinity (K_i_ = 280 nM) (Scheme 1). Strikingly, removal of the bridgehead moiety entirely abolished the activity of both compounds in luciferase reporter assays (Scheme 1), suggesting that some degree of steric occupancy is critical for the compounds’ ability to activate LRH-1. Understanding that there is a requirement for some degree of steric bulk at the bridgehead position, we searched for reaction conditions that could engage alkene **1** via conjugate addition. Because 6HP derivatives bearing heteroatom substituents at this position are known to be highly acid-sensitive,^15,26^ we targeted a protocol to forge carbon-carbon bonds. However, in line with the dearth of reactions that accept enones of this type, a broad survey of nucleophilic partners (e.g. malonates, enolates, Gillman reagents, and other organometallics) completely failed to provide the corresponding conjugate adducts.

We next turned our attention to radical coupling partners. Because the Giese reaction operates through an early transition state, these processes are less sensitive to steric hindrance.^27–29^ Drawing from our own experience in radical conjugate addition, we found that aminoalkyl radicals (readily accessed through a single electron oxidation/deprotonation sequence) readily engage olefin **1** (Fig. 3).^30,31^ More specifically, in the presence of an iridium photoredox catalyst and blue light, triethylamine was united with **1** to give rise to the corresponding adduct in 60% yield, as determined by NMR. Interestingly, these radical species appear to be uniquely effective here, as other radical sources (e.g. alkyl or aryl halides, carboxylates, NHPI esters) did not afford the desired products. This distinctive reactivity potentially owing to the special electronic properties of the α-heteroatom alkyl radical compared to aryl or unactivated alkyl radicals. Because a range of alkylamines have been demonstrated as competent coupling partners for Michael acceptors within this manifold, we presumed that this finding would grant access to a library of substituted 6HP structures.

**Fig. 3.**
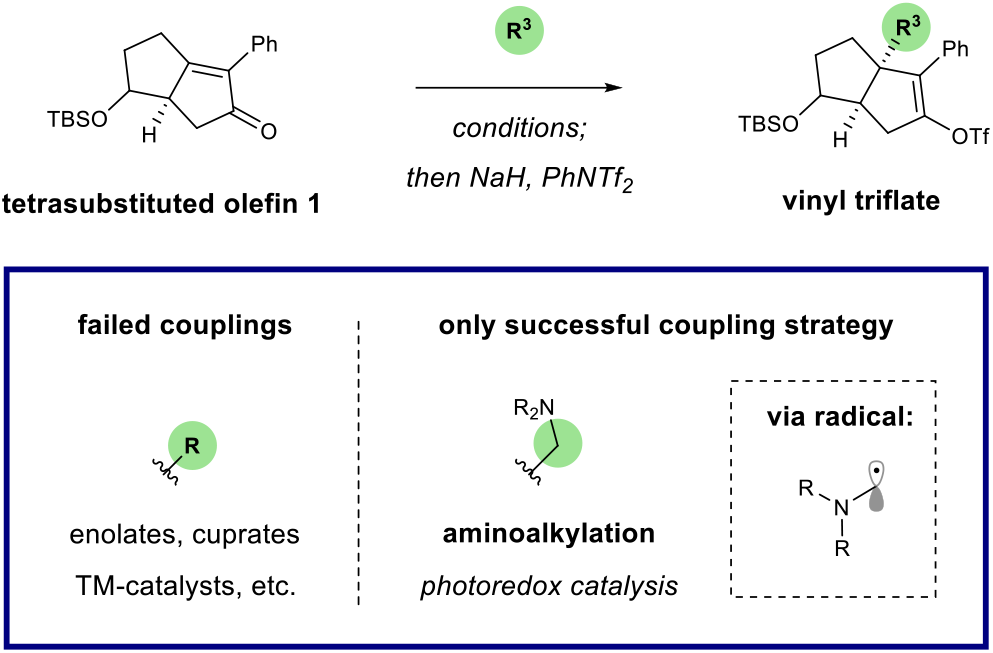
Strategy for R^3^ variation: Aminoalkyl radical conjugate addition via photoredox catalysis.

To predict the ability of 6HP derivatives with aminoalkyl substituents at the bridgehead (R^3^) position to promote binding to LRH-1, we conducted an *in silico* screen of several amine conjugate addition derivatives. Using the Glide software developed by Schrodinger,^32^ elaborated ligands derived from the Pauson-Khand product **1** (analogous to an early 6HP LRH-1 agonist, RJW100)^18^ were docked and scored. This series was conveniently selected for docking studies because we had a high-definition X-ray co-crystal structure of LRH-1 bound to the 6HP agonist RJW100 and a fragment of coregulator protein TIF-2 (PDB 5L11) to use as a reference. The scoring protocol provides XP GScores, which approximate the ΔG of binding (in kcal/mol) for each compound and these scores were used to rank the potential for each docked compound to bind to LRH-1 in a productive manner. A diverse set of cyclic and acyclic, aliphatic and aromatic, and basic and non-basic amines were docked and scored, and a selection of the results are shown in Fig. 4. Hydrophobic groups were preferred over more hydrophilic ones, with charged groups (such as protonated amines) showing a significant drop inpredicted binding affinity. Compounds derived from *N*, *N*-dialkylanilines scored the best, with the ligand derivatized from *N*,*N*-dimethylaniline scoring similarly to the parent molecule, RJW100. Overlaying the predicted binding pose of the *N*,*N*dimethylaniline-derived ligand with that of the known pose of RJW100 showed nearly perfect overlap throughout the structure, as shown in the bottom of Fig. 4. This includes in the exocyclic phenyl rings, despite the difference in linker length between the phenyl ring and bridgehead position. These data indicated that amine conjugate addition could be utilized in the design of a new class of 6HP LRH-1 agonists. To evaluate this idea, we conducted radical conjugate addition of dimethylaniline to olefin **1** under the previously outlined photon-driven reaction conditions. Again, *in situ* vinyl triflate formation gave rise to **10**, containing the completed 6HP core. As illustrated in Scheme 2, synthetic elaboration of this intermediate to the corresponding *endo*-sulfamide (**15**) and terminal carboxylate (**13**) analogs proceeded according to the previously developed protocols. Upon evaluation of these compounds using FP competition (for binding) and luciferase reporter assays (for LRH-1 transcriptional activity), we found that the bioactivity of this series essentially parallels those of the analogous bridgehead styrenes.^19,20^ Specifically, *endo*-sulfamide **15** demonstrated greater in-cell potency than the terminal carboxylate analog **13**, which presumably results from direct interactions with the polar network deep within the binding pocket (centered around the T352 hydroxyl).^19^ Further, the terminal carboxylic acid **13** showed augmented efficacy, which we propose arises from the ligand contacting phospholipid-binding residues at the mouth of the pocket.^20^

## Scheme 2.

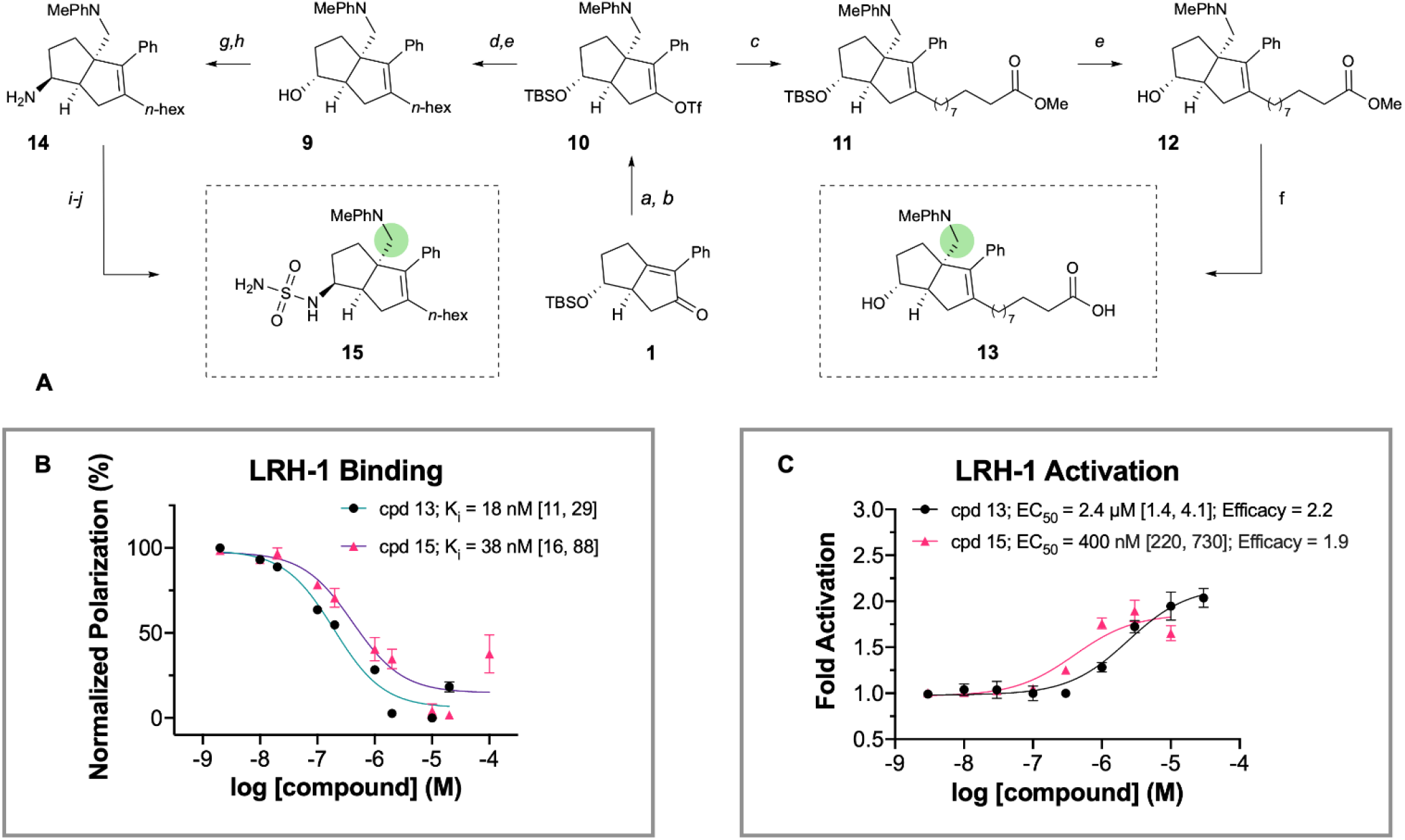
Synthesis and Evaluation of *N*,*N*-Dimethylaniline Bridgehead Compounds **A)**Reagents and conditions: (a) *N*,*N*-dimethylaniline, Ir[dF(CF_3_)ppy]_2_dtbpy·PF_6_, blue LED, 23 °C; 16 h; (b) NaH, PhNTf_2_, 0 – 23 °C; 73% yield over two steps; (c) SPhos Pd G3, SPhos, IZn(CH_2_)_9_CO_2_Me·LiCl, THF, 50 °C; 16 h; (d) SPhos Pd G3 or Pd(OAc)_2_, SPhos, IZn(CH_2_)_5_CH_3_·LiCl, THF, 50 °C; 16 h; (e) conc. aq. HCl, MeOH, 23 °C; 1 h, 49 – 54% yield over two steps; (f) LiOH, H_2_O, THF, 50 °C; 16 h, quant; (g) TPAP, NMO, MeCN, 23 °C; 1 h, 84% yield; (h) i. Ti(OiPr)_4_, NH_3_, MeOH, 23 °C; 5 h; ii. NaBH_4_, MeOH, 23 °C; 5 h, 34% yield; (i) chlorosulfonylisocyanate, *^t^*BuOH, TEA, DCM, 0 – 23 °C; 1.5 h; (j) conc. aq. HCl, dioxane, 0 – 40 °C; 14 h, 26% yield over two steps. **B)** FP evaluation of compounds 13 and 15. Data shown as mean ± SEM from two independent experiments. **C)** Luciferase reporter data for 13 and 15 shown as mean ± SEM from three biological replicates. K_i_ and EC_50_ values are given with 95% confidence intervals in brackets.

**Fig. 4.**
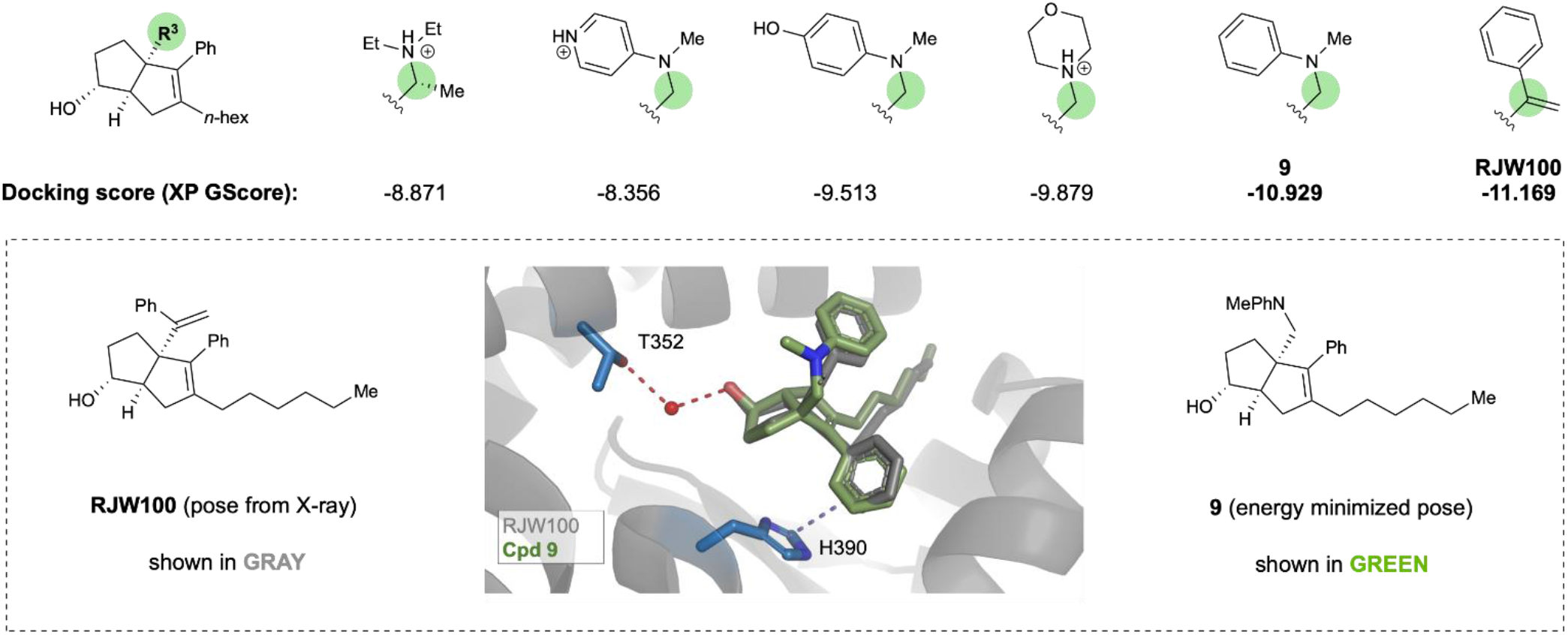
Top: The 6HP core used for docking studies and a representative sample of the screened bridgehead groups. Each group is given with the XP GScore assigned by Glide which approximates binding energy in kcal/mol. Bottom: Overlay of RJW100 (gray; PDB 5L11) and predicted minimized energy pose of **9** (green) in the LRH-1 ligand binding pocket. Key interactions with Thr352 through water and pi-stacking with His390, highlighted in blue sticks, are retained in both poses and highlighted

Encouraged by the results from FP competition and luciferase reporter assays, we assessed whether the aniline substituent promotes an active conformation at the activation function surface (AFS), which preferentially binds coregulator proteins that drive NR target gene expression.^33^ By determining how compounds drive recruitment of coregulators, we can thoroughly examine whether the aniline bridgehead group effects ligand-driven activation of LRH-1. Therefore, we used the Microarray Assay for Real-time Coregulator-Nuclear Receptor Interaction (MARCoNI), which quantifies binding of 154 peptides corresponding to NR interaction motifs from 64 coregulators with a microarray platform.^34^ We compared the coregulator binding profile of **6N**-bound and **15**-bound LRH-1 ligand-binding domain (LBD), relative to apo-LRH-1. **6N** demonstrated notable trends that involved decreased binding to corepressors, such as nuclear receptor co-repressor 1 (NCOR1) and NCOR2, and an increased binding to coactivators, such as p160/steroid receptor coactivator family member nuclear receptor coactivator 1 (NCOA1) and mediator of RNA polymerase II transcription subunit 1 (MED1). Importantly, these trends were mirrored in **15**-bound LRH-1 (Fig. 5, Fig. S1), demonstrating that the aniline substituent promotes a similar compound-mediated conformation of the AFS as the styrene bridgehead of **6N**. This shows that compounds with the aniline bridgehead group effectively drive LRH-1 activity in a similar fashion as previous agonists.

**Fig. 5.**
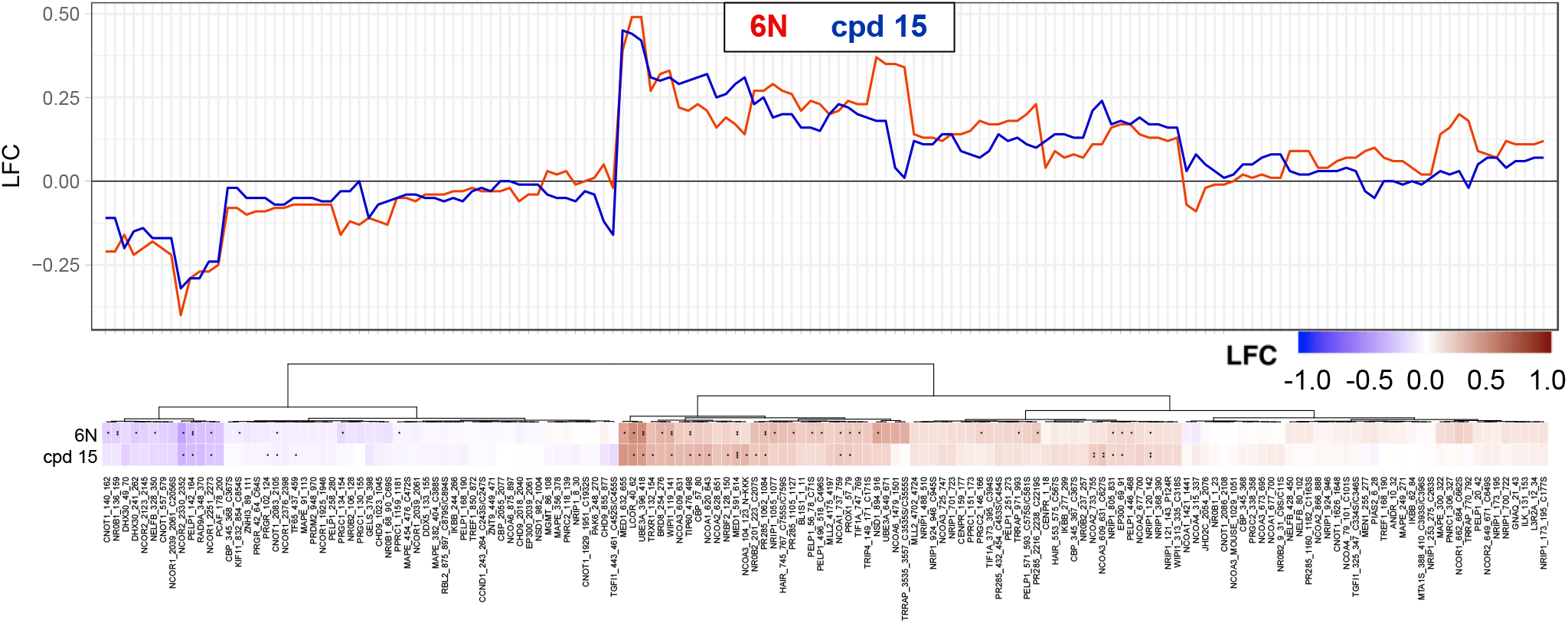
Microarray Assay for Real-time Coregulator-Nuclear Receptor Interaction (MARCoNI) comparing coregulator binding between **6N**- and **15**-bound LRH-1 LBD. Log-fold change (LFC) of peptides corresponding to the binding interface of coregulators is indicated. * p < 0.05; ** p < 0.01 – Student’s t-test, FDR.

To determine the binding pose of the aniline-containing agonists, we generated a co-crystal structure of **15** and LRH-1 (Fig. 6, Table S1). The sulfamide moiety, internal styrene, and bicyclic core of the ligand assumed the same conformation seen for other 6HP agonists (Fig. 5, bottom left).^16,19^ Surprisingly, the exocyclic aniline moiety was rotated in the opposite direction from the styrene in previous crystal structures (Fig. 6, bottom left).^16,19^ This is contrary to the prediction made by Glide, which placed the aniline phenyl group superimposed with the styrene phenyl group (Fig. 4). The observed positioning supports the hypothesis that there are no specific interactions made by the exocyclic bridgehead group and that the compound efficacy granted by its inclusion are largely a result of space-filling hydrophobic interactions. This unexpected aniline binding pose also reveals unforeseen potential for this compound series. The aniline phenyl group is oriented towards hydrophilic residues, providing novel future targets in the highly lipophilic LRH-1 binding pocket (Fig. 6, bottom right). Interestingly, these residues have been implicated in allosteric paths critical for communication between LRH-1 ligands and the AFS,^12^ suggesting that modifications targeting these residues may be an interesting route for agonist development. The LRH-1 activation-function helix (AF-H) is also within ~6 Å, providing the opportunity to directly modulate the dynamics of the AFS to induce unique gene expression profiles that may not be possible through indirect allosteric modulation.

**Fig. 6.**
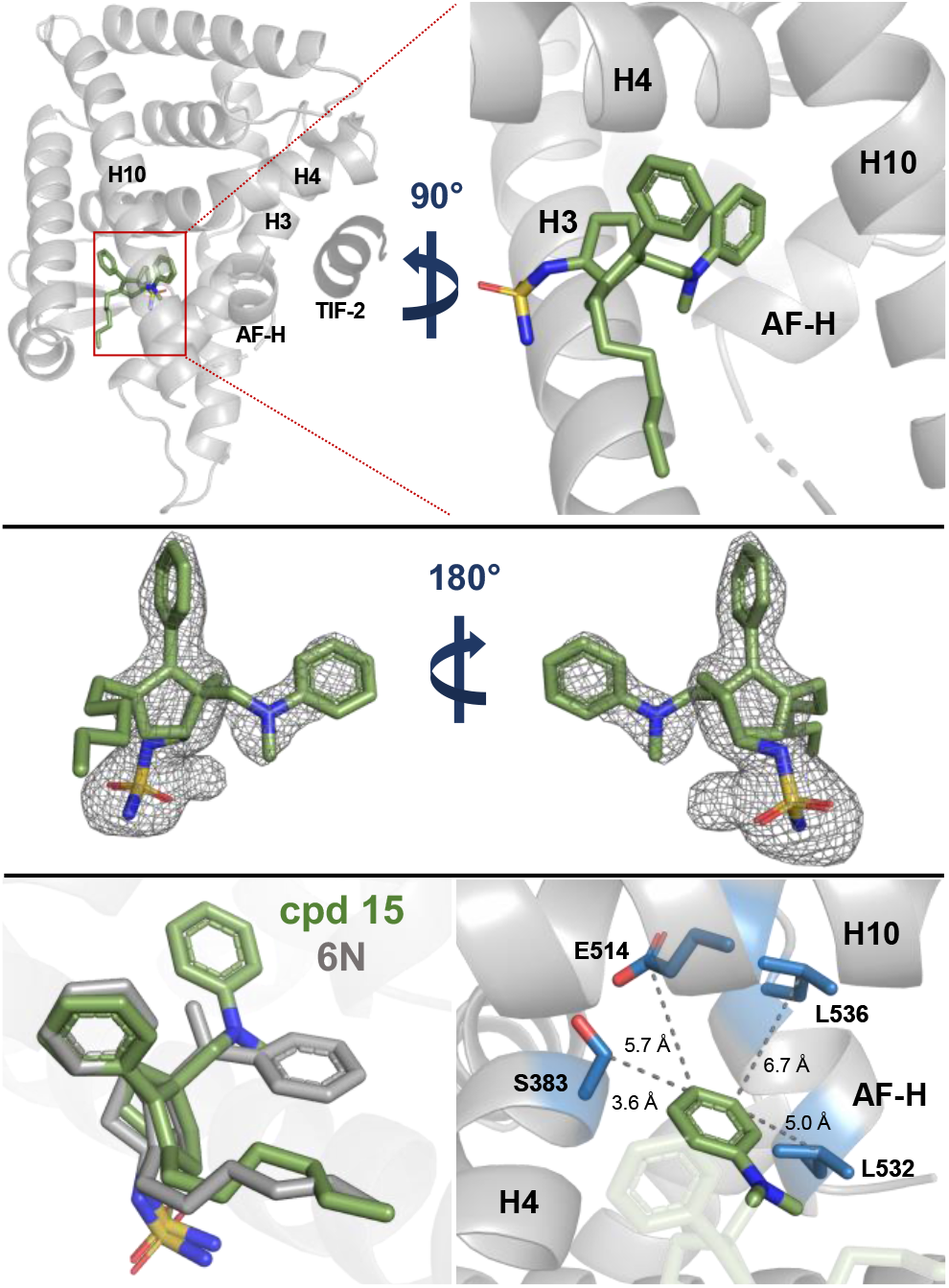
Co-crystal structure of **15** and LRH-1 reveals unexpected binding pose for the bridgehead aniline group. Top: LRH-1 (gray) and **15** (C=green, O=red, N=blue, S=yellow) in the binding pocket with the bridgehead aniline group oriented toward AF-H. Coregulator peptide fragment TIF-2 is shown in dark gray. Middle: Ligand FO-FC omit map showing electron density for **15** contoured at 2.5σ. Bottom left: **15** (green) adopts a nearly identical binding pose as **6N** (gray, PDB 6OQY), with the aniline moiety reoriented in the opposite direction from the **6N** styrene. Bottom right: Residues proximal to the aniline group are made accessible by the novel binding mode (sidechains shown as blue sticks).

In conclusion, we have developed an alternative synthesis of the standard 6HP scaffold used in modern LRH-1 agonists. This new synthesis allows for the modular modification of the bridgehead group to investigate the role of the α-styrene in 6HP LRH-1 agonists. Previous alterations to the bridgehead group, including heteroatom substitutions and small changes to the styrene, have revealed little in coregulator recruitment and luciferase reporter assays^15,19^ and have been restricted by limitations in the synthetic route. Although this group shows no clear stabilizing interactions in crystal structures,^16,19^ removal of the bridgehead group completely abolished activity in reporter assays. Guided by computational docking and enabled by photoredox, a new bridgehead moiety was installed that restored agonism while maintaining high binding affinity. A crystal structure of one of the new compounds, **15**, in the LRH-1 LBD demonstrated that the general binding pose was consistent with other 6HP agonists, though the new N,N-dimethylaniline moiety rotated to a previously unaccessed region of the binding pocket. Both the lack of critical contacts and the novel orientation of the bridgehead group suggests promise for exploitation of this novel binding mode in the development of more effective agonists and novel antagonists, which are ongoing areas of research in our laboratories.

## Acknowledgements

This research was supported by the National Institutes of Health (5R01DK115213-03). Additionally, the authors thank Cameron Pratt (Emory) for helpful discussions and suggestions related to the photoredox conjugate addition procedure.

## Appendix A. Supplementary data

Supplementary data to this article can be found online

