## Supplementary Information for "Development of a new class of liver receptor homolog-1 (LRH-1) agonists by photoredox conjugate addition"

### TABLE OF CONTENTS

**Supplementary Table 1.** X-ray data collection and refinement statistic. ……………………..2

**Supplementary Figure 1.** MARCoNI assay plots. ……………………………………………3

**Biology Methods**

*Cell Culture* ……………………………………………………………………….6
*Protein Expression* ……………………………………………………………….6

Supplementary Table 1. X-ray data collection and refinement statistics. PDB ID: 6VIF

| PDB ID: 6VIF | |
|---|---|
| **Data Collection** | |
| **Space Group** | P 43 21 2 |
| **Resolution Range (Å)** | 42.6 - 2.26 (2.34 - 2.26) |
| **Completeness (%)** | 99.7 (98.4) |
| **Unit Cell Parameters** | a = 46.13 Å, b = 46.13 Å, c = 223.44 Å<br>α = 90°, β = 90°, γ = 90° |
| **Reflections** | 21722 (2148) |
| **Unique Reflections** | 12218 (1171) |
| **Redundancy** | 16.3 (7.3) |
| **I/σ** | 17.1 (1.3) |
| **Refinement** | |
| **$R_{work}/R_{free}$** | 0.24/0.28 |
| **$R_{meas}$** | 0.217 (0.903) |
| **$R_{pim}$** | 0.050 (0.327) |
| **Ramachandran plot** | |
| **Favored (%)** | 96.7 |
| **Allowed (%)** | 3.3 |
| **Outliers (%)** | 0 |
| **B-Factors** | |
| **Protein** | 68.3 |
| **Ligand** | 67.4 |
| **Water** | 59.7 |
| **No. Non-hydrogen Atoms** | |
| **Total** | 2048 |
| **Protein** | 2004 |
| **Ligand** | 34 |
| **Water** | 10 |
| **RMS Deviations** | |
| **Bond lengths (Å)** | 0.003 |
| **Bond angles (°)** | 0.47 |

Supplementary Figure 1. Coregulator peptides in MARCoNI assay independently shown. Data represented in arbitrary fluorescence units (AU). * $p < 0.05$; ** $p < 0.01$ – Student's t-test, FDR.

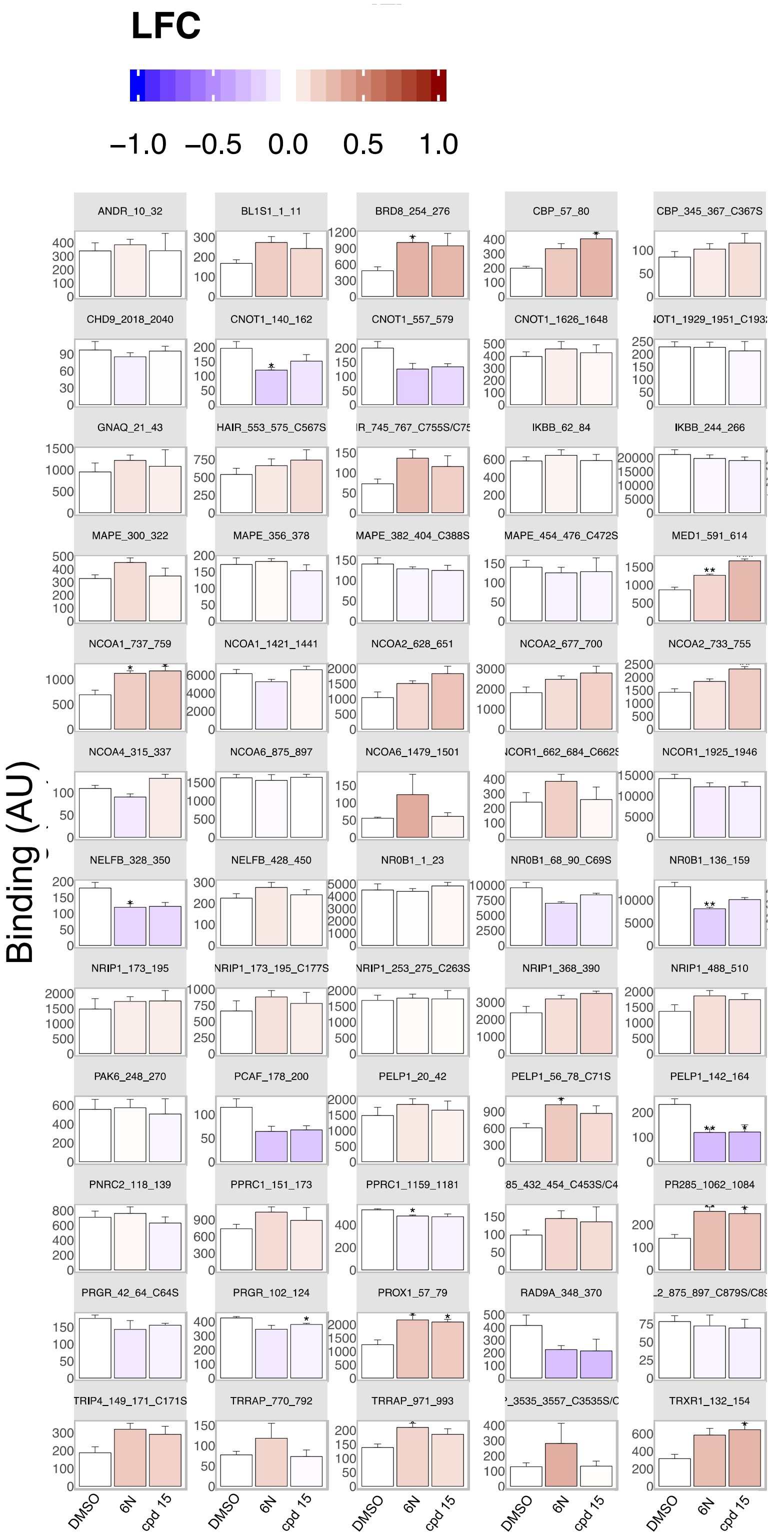

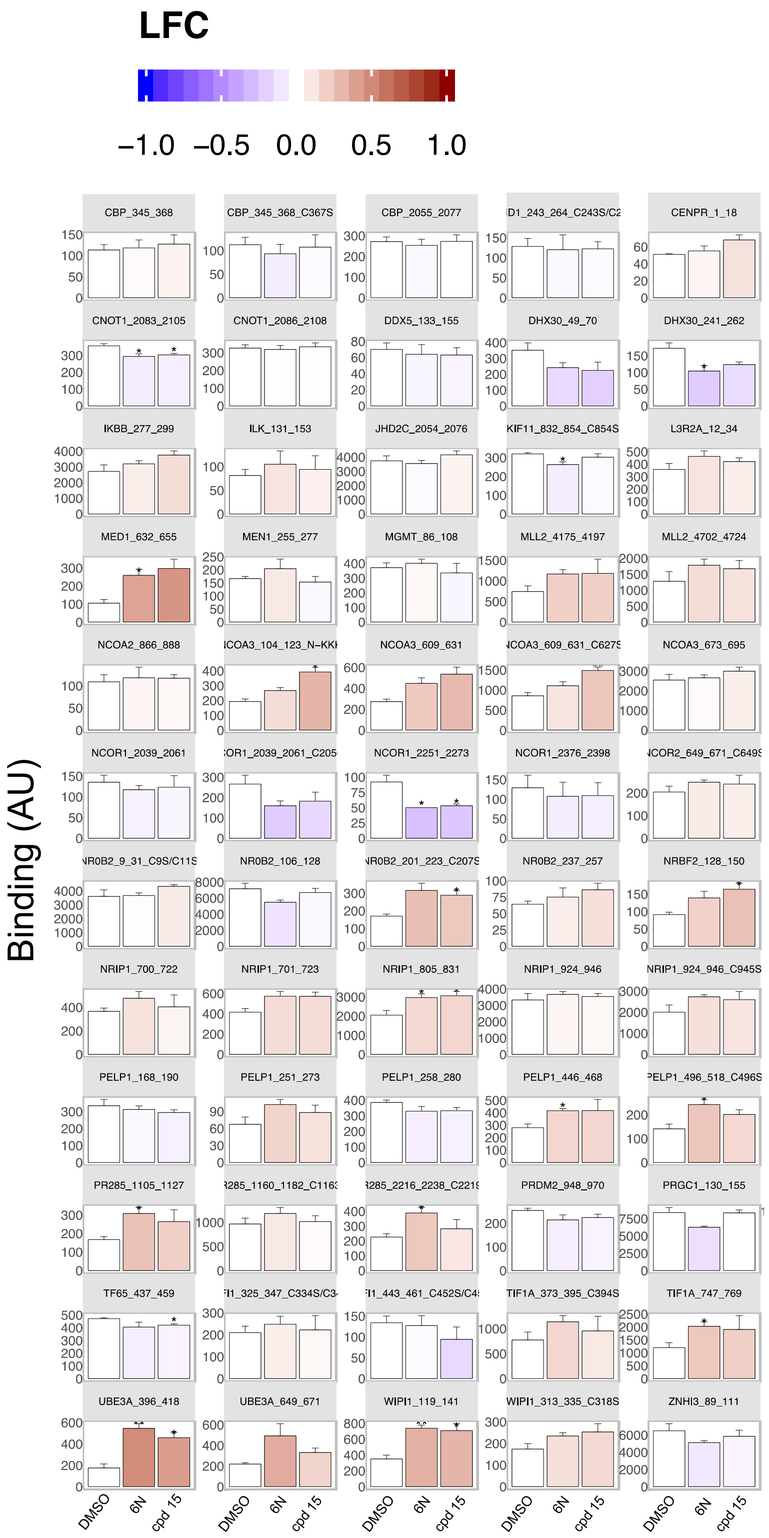
LFC
−1.0 −0.5 0.0 0.5 1.0
Binding (AU)
CBP_345_368
CBP_345_368_C367S
CBP_2055_2077
CENPR_1_18
CNOT1_2083_2105
CNOT1_2086_2108
DDX5_133_155
DHX30_49_70
DHX30_241_262
IKBB_277_299
ILK_131_153
JHD2C_2054_2076
KIF11_832_854_C854S
L3R2A_12_34
MED1_632_655
MEN1_255_277
MGMT_86_108
MLL2_4175_4197
MLL2_4702_4724
NCOA2_866_888
NCOA3_609_631
NCOA3_673_695
NCOR1_2039_2061
NCOR1_2251_2273
NCOR1_2376_2398
NR0B2_106_128
NR0B2_237_257
NRBF2_128_150
NRIP1_700_722
NRIP1_701_723
NRIP1_805_831
NRIP1_924_946
PELP1_168_190
PELP1_251_273
PELP1_258_280
PELP1_446_468
PR285_1105_1127
PRDM2_948_970
PRGC1_130_155
TF65_437_459
TIF1A_373_395_C394S
TIF1A_747_769
UBE3A_396_418
UBE3A_649_671
WIPI1_119_141
WIPI1_313_335_C318S
ZNHI3_89_111
DMSO
6N
cpd 15

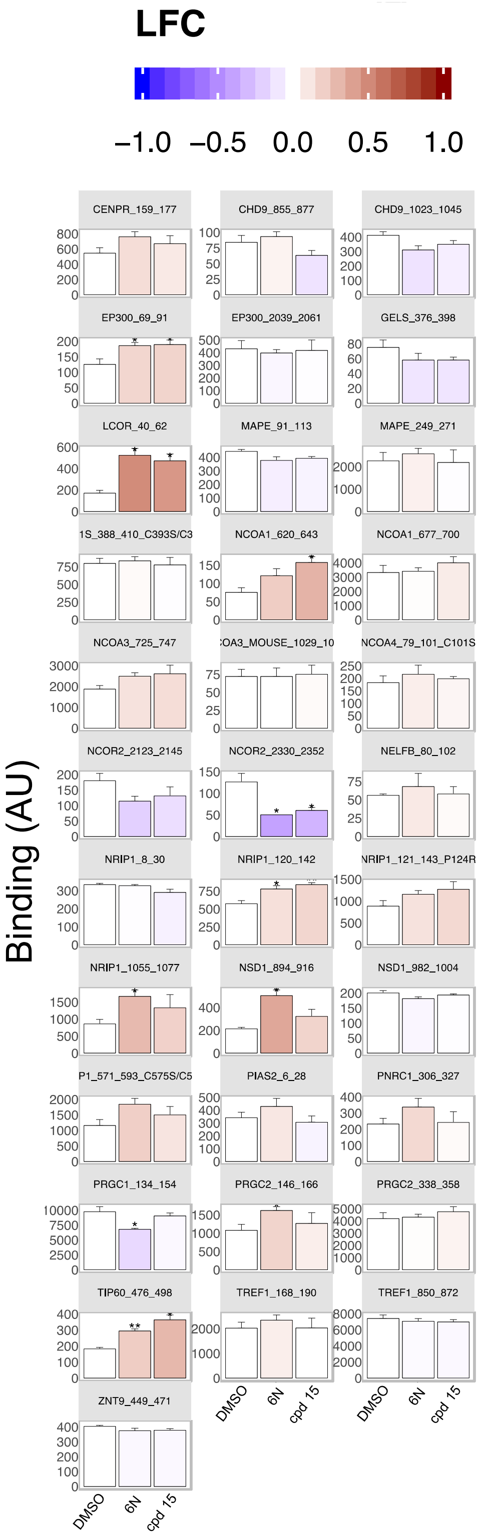
LFC
−1.0 −0.5 0.0 0.5 1.0
Binding (AU)
CENPR_159_177
CHD9_855_877
CHD9_1023_1045
EP300_69_91
EP300_2039_2061
GELS_376_398
LCOR_40_62
MAPE_91_113
MAPE_249_271
NCOA1_620_643
NCOA1_677_700
NCOA3_725_747
NCOR2_2123_2145
NCOR2_2330_2352
NELFB_80_102
NRIP1_8_30
NRIP1_120_142
NRIP1_1055_1077
NSD1_894_916
NSD1_982_1004
PIAS2_6_28
PNRC1_306_327
PRGC1_134_154
PRGC2_146_166
PRGC2_338_358
TIP60_476_498
TREF1_168_190
TREF1_850_872
ZNT9_449_471
DMSO
6N
cpd 15

### BIOLOGY METHODS

#### Cell Culture

HeLa cells were cultured in phenol red-free MEMα + 10% fetal bovine serum – Charcoal/Dextran Treated (Atlanta Biologicals) and cultured under standard conditions (5% $CO_2$, 37°C). HeLa cell identity was confirmed with an ATCC Cell Line Authentication Kit.

#### Protein Expression

BL21 pLysS *E. coli* cells were transformed with human LRH-1 LBD (residues 300-541 with N-terminal 6xHis tag) in a pMCSG7 vector and grown in liquid broth (LB) supplemented with ampicillin and chloramphenicol at 37°C. When cultures reached $OD_{600}$ 0.6, protein expression was induced with 1 mM IPTG (4 hours at 30°C). Pelleted cells were then subjected to one freeze-thaw cycle, resuspended in lysis buffer (20 mM Tris-HCl – pH 7.4, 150 mM NaCl, 5% glycerol, 25 mM imidazole, DNase, lysozyme, and PMSF), and lysed via sonication. Lysate was centrifuged (16,000 x g for 45 minutes) and the resulting supernatant was subjected to $Ni^{2+}$ affinity chromatography. Protein used for FP competition assays was incubated with DLPC (four-fold molar excess) 16 h at 4°C, purified with size-exclusion chromatography (SEC) into assay buffer (150 mM NaCl, 20 mM Tris-HCl – pH 7.4, 5 % glycerol), concentrated to ~ 3 mg/mL, and then stored at -80°C. Protein used for crystallization was incubated with TEV protease to remove the 6xHis tag, subjected to a second round of nickel affinity chromatography to separate the protein from the cleaved tag, concentrated to ~ 3.5 mg/mL, and then stored at -80°C.

#### Ligand Binding Assays

Fluorescence polarization competition assays were performed as previously described.[1] Briefly, experiments were conducted in black, polystyrene 384-well plates in assay buffer (150 mM NaCl, 20 mM Tris-HCl – pH 7.4, 5% glycerol). 6N conjugated to fluorescein amidite (FAM) (10 nM/well) was incubated with uncleaved (6xHis tag not removed) LRH-1 LBD (5 nM/well). Unlabeled competing compounds were added at concentrations indicated in figures. Each experiment was performed twice, each with four technical replicates averaged and normalized independently prior to final data analysis. GraphPad Prism (version 8) was used to analyze data, which was then fit to a one-site, fit $K_i$ curve, with a final probe concentration of 10 nM and probe affinity of 1 nM. Data was excluded from wells with 4 $e^{-4}$ M cpd 13 as the resulting point was abnormally high, presumably as a result of low solubility.

#### Reporter Assays

LRH-1 reporter assays were conducted as described previously.[2,3] Briefly, HeLa cells were seeded at 7,500 cells per well in white-walled, clear bottom 96-well plates. After ~ 24 hours, cells were transfected with LRH-1 in a pCI vector (5 ng/well), a SHP-luciferase reporter with an LRH-1 response element derived from the SHP promoter cloned upstream of firefly luciferase in a pGL3 basic vector (50 ng/well), and a *Renilla* luciferase reporter with a CMV promoter (1 ng/well). Transfection was performed using FuGENE at a ratio of 5:2 (FuGENE:DNA). Approximately 24 hours after transfection, compounds were dissolved in Opti-MEM and then introduced to cells to give final concentrations indicated in figures with DMSO at a final concentration of 0.370%. After ~ 24 hours, luciferase signal was quantified using the DualGlo kit (Promega). Each experiment was conducted with three biological replicates (corresponding to distinct passage numbers), each with three technical replicates that were averaged prior to data analysis. Firefly luciferase signal was first normalized by dividing *Renilla* signal intensity for each well and then normalizing relative to the DMSO control. Data was analyzed with GraphPad Prism (version 8) using a stimulating dose-response curve (three parameters – Hill slope = 1). Data was excluded from cells that demonstrated a high level of cell death, which was apparent from decreased size and round morphology and/or from consistently low *Renilla* signal. Using these criteria, data was excluded from analysis for cells treated with $3e^{-5}$ M of cpd 15.

**MARCoNI Assay**

*Generation of Apo LRH-1*

To generate apo LRH-1 LBD, 1 mL of purified protein (3 mg) was treated with 3.75 mL of chloroform-methanol solution (1:2 v/v) and vortexed briefly. An additional 2.5 mL chloroform:water solution (1:1 v/v) was added and the mixture was vortexed again. Protein was then pelleted by centrifugation at 1000 rpm for 10 minutes. The resulting pellet was dissolved into 0.5 mL of buffer (50 mM Tris – pH 8.0, 6 M guanidine hydrochloride, and 2 mM DTT). Protein was refolded by fast dilution at 4 °C into 25 mL of buffer (20 mM Tris – pH 8.5, 1.7 M urea, 4% glycerol and 2 mM DTT). The final urea concentration was adjusted to 2 M, and protein was concentrated to ~ 1.5 mL. Protein was then dialyzed 16 h against PBS containing 2 mM DTT at 4 °C. Refolded protein was purified by SEC to remove aggregates and unfolded protein. Refolded protein was then assessed by testing ability to bind ligand using fluorescence polarization.

*MARCoNI Assay Setup*

In vitro NR-coregulator recruitment by MARCoNI Assay mixes of 50 nM His-SUMO-hLRH-1 (apo, or preloaded with compound during purification), 25 nM ALEXA488-conjugated penta-His antibody (Qiagen # 35310), 50 µM DTT, 10 µM freshly added compound (or 2% DMSO for apo) were made in 20mM Tris (pH 7.4), 250 mM NaCl and 0.5 mM TCEP and stored on ice. LRH-1 in these assay mixes was functionally analyzed by the Microarray Assay for Real-time Coregulator Nuclear Receptor Interaction (MARCoNI), using PamChip #88101 with 154 unique coregulators sequences as described previously.[4] In short, each condition was tested using three technical replicates (arrays), and LRH-1 binding to each coregulator motif was quantified using BioNavigator software (PamGene International B.V., The Netherlands.). The modulation index (*i.e.* compound-induced log-fold change of LRH-1 binding to each coregulator) and significance of this modulation by Student's t-Test *vs*. apo LRH-1 were calculated and visualized using R software.[5] Compound and interaction (dis-)similarity were calculated by Hierarchical Clustering on Euclidean Distance and Ward's agglomeration.

**Crystallography and Structure Determination**

Complexed LRH-1 LBD crystals were generated as described previously.[2] Briefly, cleaved LRH-1 LBD (6xHis tag removed) was incubated with 6N-aniline (four-fold molar excess) for 16 h at 4°C. The complex was then purified via SEC into crystallization buffer (150 mM NaCl, 100 mM ammonium acetate, 1 mM EDTA, 2 mM CHAPS, 1 mM DTT, pH 7.4) and subsequently incubated with a peptide corresponding to human TIF-2 NR box 3 ($NH_3$-KENALLRYLLDKDD-$CO_2$) at four-fold molar excess for two hours at room temperature. The complex was then concentrated to ~ 5 mg/mL and crystals were generated via hanging drop vapor diffusion in crystallant containing 0.05 M Na acetate – pH 4.6, 5-11% PEG 4000, and 0-25% glycerol. Crystals were grown at 18°C with a microseeding approach, using LRH-1 LBD complexed with RJW100 as seed stocks (generated as described previously).[6] Crystals were then flash frozen in liquid $N_2$ using cryoprotectant consisting of crystallant supplemented with 30% glycerol. Data was collected remotely from the South East Regional Collaborative Access Team (SER-CAT) at the Advanced Photon Source (Argonne National Laboratories, Chicago, IL). Data were processed using HKL2000 and phased with molecular replacement, using PDB 6OQY (ligand omitted) as the search model. Structure refinement was performed with Phenix[7] and Coot[8], with additional refinement and assessment accomplished with PDB-REDO[9]. During refinement, residues 527-529 were removed due to poor electron density. Final figures were constructed with PyMOL.[10]

### CHEMISTRY METHODS

All reactions were carried out in oven-dried glassware, equipped with a stir bar and under a nitrogen atmosphere with dry solvents under anhydrous conditions, unless otherwise noted. Solvents used in anhydrous reactions were purified by passing over activated alumina and storing under argon. Yields refer to chromatographically and spectroscopically ($^1$H NMR) homogenous materials, unless otherwise stated. Reagents were purchased at the highest commercial quality and used without further purification, unless otherwise stated. Organic solutions were concentrated under reduced pressure on a rotary evaporator using a water bath. Chromatographic purification of products was accomplished using forced-flow chromatography on 230-400 mesh silica gel. Preparative thin-layer chromatography (PTLC) separations were carried out on 1000µm SiliCycle silica gel F-254 plates. Thin-layer chromatography (TLC) was performed on 250µm SiliCycle silica gel F-254 plates. Visualization of the developed chromatogram was performed by fluorescence quenching or by staining using $KMnO_4$, *p*-anisaldehyde, or ninhydrin stains.

$^1$H and $^{13}$C NMR spectra were obtained from the Emory University NMR facility and recorded on a Bruker Avance III HD 600 equipped with cryo-probe (600 MHz), INOVA 600 (600 MHz), INOVA 500 (500 MHz), INOVA 400 (400 MHz), VNMR 400 (400 MHz), or Mercury 300 (300 MHz), and are internally referenced to residual protio solvent signals. Data for $^1$H NMR are reported as follows: chemical shift (ppm), multiplicity (s = singlet, d = doublet, t = triplet, q = quartet, m = multiplet, dd = doublet of doublets, dt = doublet of triplets, ddd= doublet of doublet of doublets, dtd= doublet of triplet of doublets, b = broad, etc.), coupling constant (Hz), integration, and assignment, when applicable. Data for decoupled $^{13}$C NMR are reported in terms of chemical shift and multiplicity when applicable. Liquid Chromatography Mass Spectrometry (LC-MS) was performed on an Agilent 6120 mass spectrometer with an Agilent 1220 Infinity liquid chromatography inlet. Preparative High Performance Liquid chromatography (Prep-HPLC) was performed on an Agilent 1200 Infinity Series chromatograph using an Agilent Prep-C18 30 x 250 mm 10 µm column. HPLC analyses were performed using the following conditions.

Method A: A linear gradient using water and 0.1 % formic acid (FA) (Solvent A) and MeCN and 0.1% FA (Solvent B); t = 0 min, 70% B, t = 4 min, 99% B was employed on an Agilent Zorbax SB-C18 1.8 micron, 2.1 mm x 50 mm column (flow rate 0.8 mL/min). The UV detection was set to 254 nm. The LC column was maintained at ambient temperature.

Method B: An isocratic method using 65% MeCN, 45% water, and 0.1 % FA was employed on an Agilent Zorbax SB-C18 1.8 micron, 2.1 mm x 50 mm column (flow rate 0.8 mL/min). The UV detection was set to 254 nm. The LC column was maintained at ambient temperature.

Method C: A linear gradient using water and 0.1 % formic acid (FA) (Solvent A) and MeCN and 0.1% FA (Solvent B); t = 0 min, 30% B, t = 4 min, 99% B was employed on an Agilent Zorbax SB-C18 1.8 micron, 2.1 mm x 50 mm column (flow rate 0.8 mL/min). The UV detection was set to 254 nm. The LC column was maintained at ambient temperature.

Method D: An isocratic method using 95% MeCN, 5% water, and 0.1 % FA was employed on an Agilent Zorbax SB-C18 1.8 micron, 2.1 mm x 50 mm column (flow rate 0.8 mL/min). The UV detection was set to 254 nm. The LC column was maintained at ambient temperature.

Method E: A linear gradient using water and 0.1 % formic acid (FA) (Solvent A) and MeCN and 0.1% FA (Solvent B); t = 0 min, 50% B, t = 4 min, 99% B was employed on an Agilent Zorbax SB-C18 1.8 micron, 2.1 mm x 50 mm column (flow rate 0.8 mL/min). The UV detection was set to 254 nm. The LC column was maintained at ambient temperature.

Method F: A linear gradient using water and 0.1 % formic acid (FA) (Solvent A) and MeCN and 0.1% FA (Solvent B); t = 0 min, 75% B, t = 4 min, 99% B was employed on an Agilent Zorbax SB-C18 1.8 micron, 2.1 mm x 50 mm column (flow rate 0.8 mL/min). The UV detection was set to 254 nm. The LC column was maintained at ambient temperature.

Method G: An isocratic method using 75% MeCN, 25% water, and 0.1 % FA was employed on an Agilent Zorbax SB-C18 1.8 micron, 2.1 mm x 50 mm column (flow rate 0.8 mL/min). The UV detection was set to 254 nm. The LC column was maintained at ambient temperature.

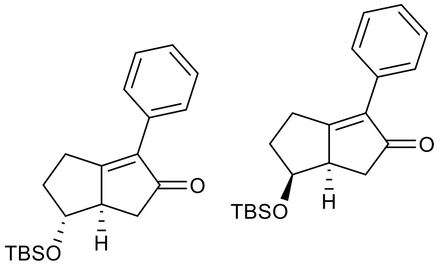

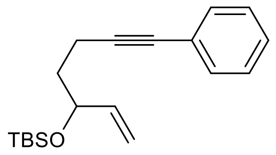

**S1**

**(6,6a)-6-((tert-butyldimethylsilyl)oxy)-3-phenyl-4,5,6,6a-tetrahydropentalen-2(1H)-one (1):** A round-bottom flask was charged with a stirbar, **tert-butyldimethyl((7-phenylhept-1-en-6-yn-3-yl)oxy)silane** (**S1**, prepared as previously reported)[3](12 mmol, 3.60 g), $Co_2(CO)_8$ (16.8 mmol, 5.74 g), and 1,2-DCE (300 ml). The resulting solution was stirred at 23 ºC while sparging with nitrogen for 3 h. The sparge was then removed and NMO (120 mmol, 14.0 g) added in small portions, using an ice bath to keep reaction approximately 23 ºC as necessary, then continued to stir at 23 ºC for 16 h. The reaction was then pushed through a plug of silica and filtrate concentrated under reduced pressure to a white solid. The crude product was purified by flash chromatography over silica with 1-10% EtOAc/hexane eluent to separate the two diastereomers, with the *exo* isomer (2.26 g) eluting first then the *endo* isomer (1.54 g) both as white solids (96% yield of combined diastereomers).

*endo* diastereomer: **$^1$H NMR** (600 MHz, $CDCl_3$) δ 7.59 (d, *J* = 8.0 Hz, 2H), 7.39 (t, *J* = 7.5 Hz, 2H), 7.30 (dd, *J* = 7.7, 0.9 Hz, 1H), 4.31 (t, *J* = 3.9 Hz, 1H), 2.99 (q, *J* = 4.8 Hz, 1H), 2.87 (dd, *J* = 18.9, 10.7 Hz, 1H), 2.80 – 2.71 (m, 1H), 2.55 (d, *J* = 5.0 Hz, 2H), 2.28 (dddd, *J* = 14.2, 10.9, 8.5, 3.7 Hz, 1H), 2.05 (ddd, *J* = 13.8, 8.3, 2.2 Hz, 1H), 0.79 (s, 9H), 0.04 (s, 3H), 0.03 (s, 3H).

**$^{13}$C NMR** (126 MHz, $CDCl_3$) δ 209.4, 183.1, 135.6, 132.2, 128.4, 128.3, 127.7, 70.6, 50.9, 37.5, 36.5, 25.9, 25.5, 18.2, -4.4, -4.8.

HPLC method C **LRMS** (ESI, APCI) m/z: calc'd for $C_{20}H_{29}O_2Si$ $(M+H)^+$ 329.2, found 328.9.

*exo* diastereomer: **$^1$H NMR** (600 MHz, $CDCl_3$) δ 7.55 (dd, J = 8.3, 1.3 Hz, 2H), 7.39 (d, J = 7.8 Hz, 2H), 7.30 (tt, J = 7.0, 1.3 Hz, 1H), 3.78 (td, J = 9.2, 7.3 Hz, 1H), 3.10 – 3.00 (m, 2H), 2.81 (dd, J = 18.0, 6.4 Hz, 1H), 2.70 – 2.60 (m, 1H), 2.33 (dd, J = 18.5, 2.8 Hz, 1H), 2.29 – 2.20 (m, 1H), 2.13 – 2.03 (m, 1H), 0.91 (s, 9H), 0.08 (s, 3H), 0.08 (d, J = 534.1 Hz, 3H).

**$^{13}$C NMR** (126 MHz, $CDCl_3$) δ 207.9, 180.5, 136.3, 131.4, 128.5, 128.4, 128.1, 77.6, 51.8, 41.6, 35.3, 26.5, 26.0, 18.2, -4.4, -4.5.

HPLC method C **LRMS** (ESI, APCI) m/z: calc'd for $C_{20}H_{29}O_2Si$ $(M+H)^+$ 329.2, found 328.9.

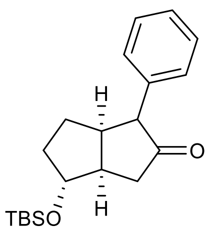

**(3a,4,6a)-4-((tert-butyldimethylsilyl)oxy)-1-phenylhexahydropentalen-2(1H)-one (S2):** A round-bottom flask was charged with a stirbar, **1** (2.55 mmol, 839.3 mg), and palladium on carbon (2.5 mol%, 272.3 mg) and then evacuated and backfilled with nitrogen four times. Dry toluene (15 mL) and acetic acid (5.12 mmol, 293 µL) were added to the reaction flask and allowed to stir at 23 °C. The reaction flask was opened briefly and $NaBH_4$ (5.12 mmol, 193.6 mg) was added under positive pressure. The reaction was allowed to stir for 1 h and then quenched with 0.1 M HCl until bubbling ceased before exposing to the atmosphere. The reaction solution was made basic using saturated $NaHCO_3$ solution and quickly extracted two times with EtOAc. The resultant organic layers were dried over $Na_2SO_4$ and filtered through Celite. Filtrate was concentrated under reduced vacuum to produce a milky oil which was immediately evacuated and backfilled with nitrogen four times to avoid decomposition in air. The oil was then dissolved in dry benzene under nitrogen and was used without further purification in subsequent steps.

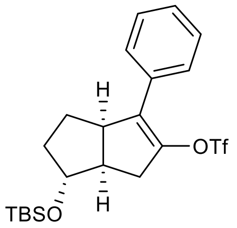

**(3a,6,6a)-6-((tert-butyldimethylsilyl)oxy)-3-phenyl-1,3a,4,5,6,6a-hexahydropentalen-2-yl trifluoromethanesulfonate (2):** A flame-dried round-bottom flask was charged with a stirbar and NaH (60% dispersion in mineral oil, 5.1 mmol, 204 mg) then evacuated and backfilled with nitrogen four times. Dry DMF (26 ml) was then added, and the reaction flask was cooled to 0 ºC. **S2** (approximately 2.55 mmol) was added slowly as a solution in dry benzene via syringe. After stirring at 0 ºC for 2 h, $PhNTf_2$ (3.83 mmol, 1.366g) was added as a solid and reaction put back under nitrogen. The resulting mixture was allowed to warm to 23 ºC and stirred 16 h. The mixture was then quenched with EtOAc before exposing to atmosphere and further diluting with EtOAc and $H_2O$. The organic layer was washed four times with $H_2O$ then brine, dried over $Na_2SO_4$, and filtered. Filtrate was concentrated under reduced pressure to obtain a brown oil. The crude product was purified by flash chromatography on silica with EtOAc/hexane eluent (1-10%) to obtain the title compound as a clear oil (817 mg, 69% over 2 steps from conjugate reduction).

**$^1$H NMR** (400 MHz, $CDCl_3$) δ 7.43 (d, J = 7.7 Hz, 2H), 7.36 (t, J = 7.2 Hz, 2H), 7.30 (t, J = 7.2 Hz, 1H), 3.97 (q, J = 3.8 Hz, 1H), 3.70 (t, J = 8.6 Hz, 1H), 3.07 (dd, J = 17.1, 10.2 Hz, 1H), 2.62 (t, J = 9.8 Hz, 1H), 2.49 (dt, J = 17.0, 3.6 Hz, 1H), 2.11 – 1.99 (m, 1H), 1.73 – 1.51 (m, 2H), 1.43 – 1.35 (m, 1H), 0.87 (s, 9H), 0.06 (s, 3H), 0.05 (s, 3H).

**$^{13}$C NMR** (126 MHz, $CDCl_3$) δ 140.1, 132.5, 131.9, 128.5, 128.3, 128.0, 118.3 (q, *J* = 320.4 Hz), 80.3, 46.8, 45.7, 36.4, 33.5, 28.1, 25.8, 18.0, -4.6, -4.8.

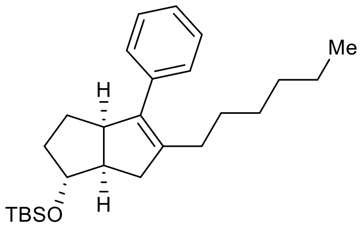

**tert-butyl(((1,3a,6a)-5-hexyl-4-phenyl-1,2,3,3a,6,6a-hexahydropentalen-1-yl)oxy)dimethylsilane (S3):** A round-bottom flask was charged with a stirbar and LiCl (15 mmol, 636 mg) then heated to 140 ºC under vacuum for 10 minutes before cooling again to 23 ºC. Once cooled, zinc (30 mesh, 22.5 mmol, 1.47 g) was added and re-heated to 140 ºC under vacuum for 10 minutes. While cooling back to 23 ºC, the flask was backfilled with nitrogen and evacuated three times. Once the flask cooled, dry THF was added (15 ml) and began stirring vigorously. To the vigorously stirred suspension was added 1,2-dibromoethane (0.75 mmol, 60 µl), trimethylsilyl chloride (0.15 mmol, 10.5 µl), and two drops of a 1M solution of $I_2$ in dry THF under nitrogen. Once the yellow color of the $I_2$ had disappeared (about 10 minutes), 1-iodohexane (15 mmol, 2.21 ml) was added neat via syringe and the solution was heated to reflux for 10 seconds then to 50 ºC. After stirring at 50 ºC for 4 h, a titer for the hexylzinc iodide of 0.50 M was obtained by colorimetric titration of an aliquot with a 1M solution of $I_2$ in dry THF (equivalence point reached when $I_2$ color persists with stirring). A separate flame-dried reaction vial was charged with a stirbar, **2** (0.108 mmol, 50 mg), SPhos G3 (5.4 µmol, 4.2 mg), and SPhos (10.8 µmol, 4.4 mg). The reaction vial was evacuated and backfilled with nitrogen four times then dry THF (0.3 ml) added and the resulting solution stirred at 50 ºC. After 5 minutes hexylzinc iodide solution added (0.324 mmol, 0.648 ml) via syringe. The resulting mixture was heated to 50 ºC for 16 h before cooling back to 23 ºC and pushing through a plug of silica with ethyl acetate. Filtrate concentrated under reduced pressure to a black oil. The crude product was used in subsequent steps without further purification.

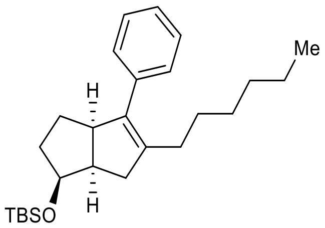

**(1,3a,6a)-5-hexyl-4-phenyl-1,2,3,3a,6,6a-hexahydropentalen-1-ol (6):** A round-bottom flask was charged with a stirbar and **S3** (approximately 0.108 mmol). The material was suspended in MeOH (2 ml) and DCM was added until all of **S3** had dissolved. The resulting solution was stirred at 23 ºC and two drops of concentrated hydrochloric acid added. After 1 h the reaction was diluted with EtOAc and washed with saturated aqueous $NaHCO_3$, $H_2O$ twice, then brine. The organic layer was dried over $Na_2SO_4$, filtered and concentrated under reduced pressure to collect a crude mixture. The crude mixture was purified by flash chromatography over silica with 10-30% EtOAc/hexanes eluent to collect the title compound (7.5 mg, 24% yield over 2 steps).

**$^{1}$H NMR** (500 MHz, $CDCl_3$) δ 7.33 (t, J = 7.4 Hz, 2H), 7.23 (dd, J = 7.4, 1.4 Hz, 1H), 7.20 – 7.17 (m, 2H), 4.02 (q, J = 3.5 Hz, 1H), 3.73 (t, J = 8.3 Hz, 1H), 2.77 (dd, J = 17.2, 10.0 Hz, 1H), 2.59 (t, J = 9.2 Hz, 1H), 2.26 (dt, J = 17.1, 3.2 Hz, 1H), 2.19 – 2.09 (m, 1H), 2.09 – 2.00 (m, 1H), 1.91 – 1.82 (m, 1H), 1.69 – 1.61 (m, 1H), 1.61 – 1.54 (m, 1H), 1.48 – 1.33 (m, 3H), 1.31 – 1.18 (m, 6H), 0.87 (t, J = 7.0 Hz, 3H).

**$^{13}$C NMR** (126 MHz, $CDCl_3$) δ 138.3, 138.2, 137.7, 128.5, 128.0, 126.2, 81.3, 53.3, 48.3, 41.2, 33.4, 31.7, 29.3, 29.2, 28.2, 27.8, 22.6, 14.1.m

HPLC method A **LRMS** (ESI, APCI) m/z: calc'd for $C_{20}H_{27}$ $(M\text{-}OH)^+$ 267.2, found 267.0.

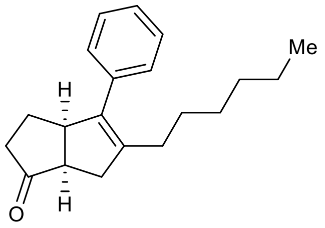

**(3a,6a)-5-hexyl-4-phenyl-3,3a,6,6a-tetrahydropentalen-1(2H)-one (S4):** A reaction vial was charged with a stirbar, **6** (0.574 mmol, 163.2 mg), and MeCN (5.7 ml). The resulting solution stirred at 23 ºC then TPAP (57 µmol, 20.2 mg) and NMO (5.73 mmol, 672.2 mg) added. The reaction solution continued to stir until **6** consumed by TLC before eluting through a plug of silica. The resulting crude material was then loaded on silica and eluted with 5-10% EtOAc/hexanes to collect the title compound (130.8 mg, 81% yield).

**$^1$H NMR** (500 MHz, $CDCl_3$) δ 7.36 (t, J = 7.7 Hz, 2H), 7.26 (tt, J = 6.8, 1.3 Hz, 1H), 7.16 (d, J = 8.1, 1.1 Hz, 2H), 3.96 – 3.91 (m, 1H), 2.78 – 2.62 (m, 3H), 2.24 – 1.89 (m, 5H), 1.87 – 1.81 (m, 1H), 1.43 – 1.31 (m, 2H), 1.29 – 1.13 (m, 6H), 0.85 (t, J = 7.1 Hz, 3H).

**$^{13}$C NMR** (126 MHz, $CDCl_3$) δ 224.2, 141.1, 137.3, 137.0, 128.2,3126.6, 50.9, 48.8, 39.4, 36.1, 31.6, 29.3, 29.2, 27.9, 23.9, 22.6, 14.0.

HPLC method A **LRMS** (ESI, APCI) m/z: calc'd for $C_{20}H_{26}O$ $(M+H)^+$ 283.2, found 283.0.

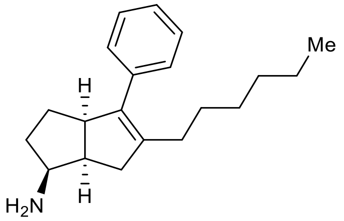

**(1,3a,6a)-5-hexyl-4-phenyl-1,2,3,3a,6,6a-hexahydropentalen-1-amine (7):** A reaction vial was charged with a stirbar, **S4** (0.463 mmol, 130.8 mg), $Ti(O^iPr)_4$ (0.694 mmol, 211 µl), and EtOH (4.6 ml) and then sealed. A solution of $NH_3$ in MeOH (7N, 9.26 mmol, 1.323 ml) was then injected and the resulting solution was stirred at 23 ºC for 6 h before unsealing vial and adding $NaBH_4$ (1.389 mmol, 52.5 mg) and continuing stirring at 23 ºC for 16 h. Reaction was then diluted with EtOAc and saturated aqueous Rochelle's salt and sonicated for 5 min. The resulting slurry was washed two times with saturated aqueous Rochelle's salt, $H_2O$, then brine. The organic layer was dried over $Na_2SO_4$, filtered, and filtrate concentrated under reduced pressure to collect the crude material. Crude material purified by flash chromatography on silica with 5:95:0 to 30:69:1 EtOAc:hexanes:$Et_3N$ eluent to collect the title compound as a single diastereomer (99.5 mg, 59% yield).

**$^1$H NMR** (500 MHz, $CDCl_3$) δ 7.32 (t, J = 7.6 Hz, 2H), 7.23 – 7.17 (m, 3H), 3.59 (t, J = 8.8 Hz, 1H), 3.36 – 3.28 (m, 1H), 2.80 – 2.72 (m, 1H), 2.58 (dt, J = 17.3, 3.5 Hz, 1H), 2.42 (dd, J = 17.3, 9.8 Hz, 1H), 2.21 – 2.13 (m, 1H), 2.13 – 2.04 (m, 1H), 1.71 – 1.64 (m, 1H), 1.60 – 1.46 (m, 1H), 1.46 – 1.32 (m, 2H), 1.33 – 1.17 (m, 9H), 0.86 (t, J = 6.9 Hz, 3H).

**$^{13}$C NMR** (126 MHz, $CDCl_3$) δ 139.2, 138.6, 138.2, 128.5, 127.9, 126.1, 55.9, 53.9, 43.1, 35.8, 33.8, 31.7, 29.3, 28.6, 28.3, 22.6, 14.1.

HPLC method A **LRMS** (ESI, APCI) m/z: calc'd for $C_{20}H_{29}N$ $(M+H)^+$ 284.2, found 284.0.

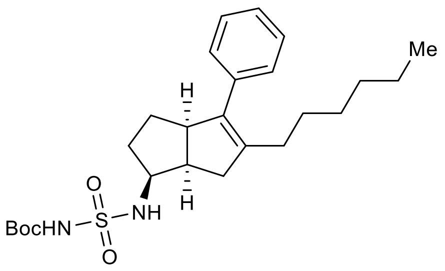

**tert-butyl (N-((1,3a,6a)-5-hexyl-4-phenyl-1,2,3,3a,6,6a-hexahydropentalen-1-yl)sulfamoyl)carbamate (S5):** An oven-dried vial was charged with a stirbar, $^t$BuOH (1.23 mmol, 91.7 mg), and DCM (12.5 ml) then evacuated under reduced pressure and backfilled with nitrogen three times and cooled to 0 ºC. Chlorosulfonyl isocyanate (1.125 mmol, 97 µl) was then added dropwise via syringe and the solution allowed to warm to 23 ºC over 90 minutes. A 2.64 ml portion of this solution was added slowly via syringe to a solution of **7** (0.225 mmol, 63.9 mg) and $Et_3N$ (0.451 mmol, 63 µl) in DCM (2.25 ml) at 0 ºC under nitrogen. This combined solution was allowed to warm to 23 ºC gradually 16 h then diluted with EtOAc. The diluted solution was washed with three times with $NH_4Cl$ then $H_2O$ and brine. The organic layer was dried over $Na_2SO_4$, filtered, and filtrate concentrated under reduced pressure to collect the crude material. Crude material purified by flash chromatography on silica with 10:90:0 to 50:49:1 EtOAc:hexanes:$Et_3N$ to give the title compound (32.7 mg, 31% yield).

**$^1$H NMR** (500 MHz, $CDCl_3$) δ 7.33 (t, J = 7.4 Hz, 2H), 7.23 (t, J = 7.3 Hz, 1H), 7.19 – 7.15 (m, 2H), 5.18 (d, J = 7.6 Hz, 1H), 3.76 (dtd, J = 9.7, 7.7, 5.6 Hz, 1H), 3.61 (t, J = 8.7 Hz, 1H), 2.95 (qd, J = 7.9, 5.7 Hz, 1H), 2.55 (d, J = 6.1 Hz, 2H), 2.21 – 2.13 (m, 1H), 2.13 – 2.04 (m, 1H), 1.83 – 1.75 (m, 1H), 1.69 – 1.50 (m, 2H), 1.50 (s, 9H), 1.49 – 1.34 (m, 2H), 1.33 – 1.17 (m, 7H), 0.86 (t, J = 7.0 Hz, 3H).

**$^{13}$C NMR** (126 MHz, $CDCl_3$) δ 150.1, 139.4, 138.1, 137.5, 128.4, 128.1, 126.4, 83.7, 58.4, 52.9, 41.5, 36.9, 31.6, 30.2, 29.3, 29.2, 28.2, 28.0, 27.9, 22.6, 14.1.

HPLC method A **LRMS** (ESI, APCI) m/z: calc'd for $C_{21}H_{30}N_2O_4S$ $(M-C_4H_8)^+$ 406.2, found 406.8.

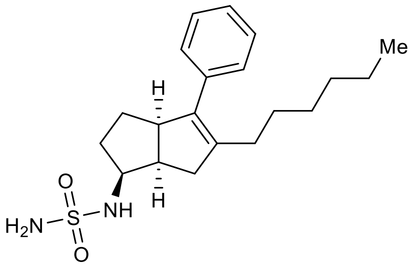

**N-((1)-5-hexyl-4-phenyl-1,2,3,3a,6,6a-hexahydropentalen-1-yl)sulfamide (8):** A reaction vial was charged with a stirbar, **S5** (70 µmol, 32.7 mg), and dioxane (530 µL). The solution was frozen in an ice bath and then allowed to slowly warm to 23 ºC. As soon was the entire solution had re-melted, cold concentrated HCl (176 µL) was added so the solution was 3:1 HCl: Dioxane. The solution was allowed to slowly warm to 23 ºC and continue reacting at 40 °C until **S5** was consumed. The reaction solution was diluted with EtOAc and washed four times with $H_2O$ then twice with brine. The organic layer was dried over $Na_2SO_4$, filtered, and filtrate concentrated under reduced pressure to collect the crude material. This crude material was purified by flash chromatography on silica with 10-40% EtOAc/hexanes to collect the title compound (19.8 mg, 77% yield).

$^{1}$**H NMR** (600 MHz, $CDCl_3$) δ 7.30 (d, J = 7.3 Hz, 2H), 7.20 (tt, J = 7.2, 1.3 Hz, 1H), 7.16 – 7.13 (m, 2H), 4.53 (s, 2H), 4.44 (d, J = 7.7 Hz, 1H), 3.84 – 3.77 (m, 1H), 3.60 (t, J = 8.4 Hz, 1H), 2.96 (dddd, J = 8.4, 4.6, 0.4 Hz, 1H), 2.57 – 2.45 (m, 2H), 2.18 – 2.10 (m, 1H), 2.10 – 2.02 (m, 1H), 1.85 – 1.79 (m, 1H), 1.63 – 1.55 (m, 2H), 1.49 – 1.34 (m, 3H), 1.28 – 1.16 (m, 6H), 0.83 (t, J = 7.1 Hz, 3H).

$^{13}$**C NMR** (126 MHz, $CDCl_3$) δ 139.2, 138.3, 137.5, 128.4, 128.1, 126.4, 57.9, 53.0, 41.3, 36.9, 31.6, 30.9, 29.3, 29.3, 28.3, 27.9, 22.61, 14.1.

HPLC method A **LRMS** (ESI, APCI) m/z: calc'd for $C_{25}H_{30}N_2O_2S$ $(M+H)^+$ 363.2, found 362.9.

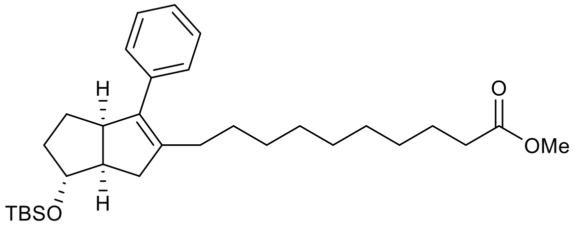

**methyl 10-((3a,6,6a)-6-((tert-butyldimethylsilyl)oxy)-3-phenyl-1,3a,4,5,6,6a-hexahydropentalen-2-yl)decanoate (3):** A flame-dried reaction vial was charged with a stirbar, **2** (0.700 mmol, 323.8 mg), Sphos G3 (35 µmol, 27.3 mg), and SPhos (70 µmol, 28.7 mg). The reaction vial was evacuated and backfilled with nitrogen four times then THF (2.3 ml) added and began heating to 50 ºC. After 10 minutes, a previously prepared alkylzinciodide solution was added (0.7 M, 2.1 mmol, 3 ml) via syringe. The resulting mixture continued to stir at 50 ºC 16 h before cooling back to 23 ºC and pushing through a silica plug with ethyl acetate. The crude product was carried on to subsequent steps without further purification.

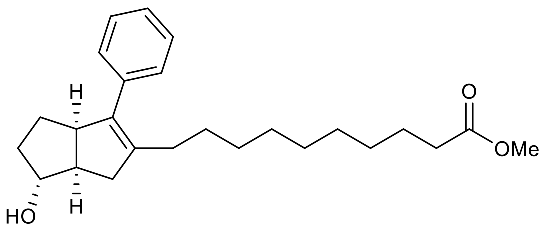

**methyl 10-((3a,6,6a)-6-hydroxy-3-phenyl-1,3a,4,5,6,6a-hexahydropentalen-2-yl)decanoate (4):** A round-bottom flask was charged with a stirbar and **3** (0.311 mmol, 155.1 mg). The material was suspended in MeOH (10 ml) and DCM was added until it dissolved. The resulting solution was stirred at 23 ºC and two drops of concentrated hydrochloric acid added. After stirring 16 h, the reaction was diluted with EtOAc and washed with saturated aqueous NaHCO3, H2O twice, then brine. The organic layer was dried over Na2SO4, filtered and concentrated under reduced pressure to collect a crude mixture. The crude mixture was purified by flash chromatography over silica with 10-30% EtOAc/hexanes eluent to collect the title compound (106.3 mg, 41% yield over 2 steps from Negishi coupling).

**$^1$H NMR** (500 MHz, $CDCl_3$) δ 7.32 (t, J = 7.5 Hz, 2H), 7.22 (t, J = 7.4 Hz, 1H), 7.18 (d, J = 7.8 Hz, 2H), 4.01 (q, J = 3.3 Hz, 1H), 3.72 (t, J = 8.8 Hz, 1H), 3.67 (s, 3H), 2.76 (dd, J = 17.1, 9.9 Hz, 1H), 2.58 (t, J = 9.2 Hz, 1H), 2.30 (t, J = 7.6 Hz, 2H), 2.25 (dt, J = 17.1, 3.1 Hz, 1H), 2.17 – 2.09 (m, 1H), 2.08 – 2.00 (m, 1H), 1.90 – 1.81 (m, 1H), 1.69 – 1.52 (m, 4H), 1.44 – 1.32 (m, 2H), 1.32 – 1.17 (m, 11H).

**$^{13}$C NMR** (126 MHz, $CDCl_3$) δ 174.3, 138.3, 138.2, 137.7, 128.4, 128.0, 126.2, 81.3, 53.3, 51.4, 48.4, 41.1, 34.1, 33.4, 29.5, 29.4, 29.3, 29.2, 29.1, 28.2, 27.8, 24.9.

HPLC method A **LRMS** (ESI, APCI) m/z: calc'd for $C_{25}H_{35}O_2$ $(M\text{-}OH)^+$ 367.3, found 366.9.

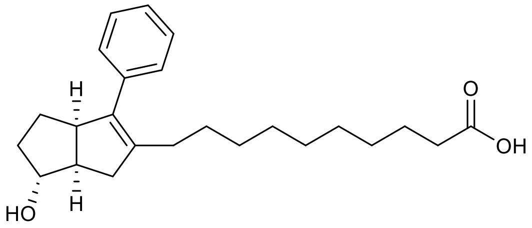

**10-((3a,6,6a)-6-hydroxy-3-phenyl-1,3a,4,5,6,6a-hexahydropentalen-2-yl)decanoic acid (5):** A reaction vial was charged with a stirbar, **4** (0.287 mmol, 106.3 mg), LiOH•$H_2O$ (2.87 mmol, 68.7 mg), and 2 ml of 5:1 THF/$H_2O$ solution. The resulting suspension was stirred at 50 ºC 16 h. The reaction was then acidified with 1 M HCl, diluted with EtOAc and $H_2O$. The aqueous layer was extracted three times with EtOAc and the organic layers were combined, washed twice with brine, dried over $Na_2SO_4$, filtered and concentrated under reduced pressure to afford the title compound (100 mg, 97% yield).

**$^1$H NMR** (600 MHz, $CDCl_3$) δ 7.29 (t, J = 7.6 Hz, 2H), 7.19 (tt, J = 7.4, 1.5 Hz, 1H), 7.16 – 7.13 (m, 2H), 3.99 (q, J = 3.6 Hz, 1H), 3.69 (t, J = 10.6 Hz, 1H), 2.73 (dd, J = 16.9, 9.8 Hz, 1H), 2.55 (t, J = 9.2 Hz, 1H), 2.32 (t, J = 7.5 Hz, 2H), 2.22 (dt, J = 17.1, 3.2 Hz, 1H), 2.13 – 2.06 (m, 1H), 2.06 – 1.97 (m, 1H), 1.86 – 1.78 (m, 1H), 1.65 – 1.57 (m, 3H), 1.57 – 1.51 (m, 1H), 1.41 – 1.31 (m, 1H), 1.32 – 1.27 (m, 1H), 1.28 – 1.09 (m, 11H).

**$^{13}$C NMR** (126 MHz, $CDCl_3$) δ 179.4, 138.26, 138.25, 137.6, 128.4, 128.0, 126.2, 81.4, 53.3, 48.3, 41.1, 34.0, 33.3, 29.5, 29.3, 29.3, 29.2, 29.1, 29.0, 28.1, 27.8, 24.7.

HPLC method A **LRMS** (ESI, APCI) m/z: calc'd for $C_{24}H_{33}O_2$ $(M\text{-}OH)^+$ 353.2, found 353.0.

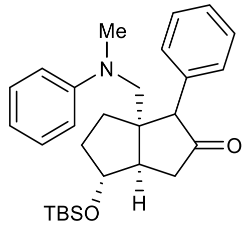

**(3a,4,6a)-4-((tert-butyldimethylsilyl)oxy)-6a-((methyl(phenyl)amino)methyl)-1-phenylhexahydropentalen-2(1H)-one (S6):** A flame-dried round-bottom flask was charged with a stirbar, **1** (0.52 mmol, 170.0 mg), and (Ir[dF($CF_3$)ppy]$_2$dtbbpy)$PF_6$ (1.33 µmol, 1.5 mg) then placed under vacuum. The flask was then evacuated and backfilled with nitrogen four times before adding freshly distilled and degassed *N*,*N*-dimethylaniline (5.2 ml) via canula. The reaction was then stirred at 23 ºC under nitrogen with blue LED lamp irradiation until starting material was consumed as monitored by $^1$H NMR of small aliquots after 22 h. The light was then turned off and the round-bottom fitted with a distillation head and placed under vacuum, being careful to keep exposure to ambient atmosphere to a minimum. The reaction flask under distillation head was slowly warmed to 80 ºC under vacuum and this temperature was maintained until all *N*,*N*-dimethylaniline residue evaporated from the flask. The distillation head was then removed and the round-bottom flask quickly capped with a septum and evacuated then backfilled with nitrogen four times. The resulting thick oil was dissolved in dry benzene under nitrogen and frozen to be used in subsequent steps without further purification.

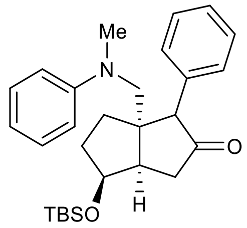

**(3a,4,6a)-4-((tert-butyldimethylsilyl)oxy)-6a-((methyl(phenyl)amino)methyl)-1-phenylhexahydropentalen-2(1H)-one (S7):** A flame-dried round-bottom flask was charged with a stirbar, **1** (2.0 mmol, 657.8 mg), and (Ir[dF($CF_3$)ppy]$_2$dtbbpy)$PF_6$ (2.0 µmol, 2.2 mg) then placed under vacuum. The flask was then evacuated and backfilled with nitrogen four times before adding freshly distilled and degassed *N*,*N*-dimethylaniline (20 ml) via canula. The reaction was then stirred at 23 ºC under nitrogen with blue LED lamp irradiation until starting material was consumed as monitored by $^1$H NMR of small aliquots after 14 h. The light was then turned off and the round-bottom fitted with a distillation head and placed under vacuum, being careful to keep exposure to ambient atmosphere to a minimum. The reaction flask under distillation head was slowly warmed to 80 ºC under vacuum and this temperature was maintained until all *N*,*N*-dimethylaniline residue evaporated from the flask. The distillation head was then removed, and the round-bottom flask quickly capped with a septum and evacuated then backfilled with nitrogen four times. The resulting thick oil was dissolved in dry benzene under nitrogen and frozen to be used in subsequent steps without further purification.

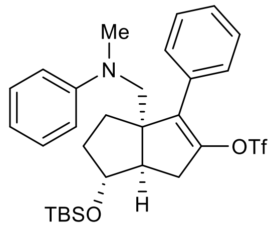

**(3a,6,6a)-6-((tert-butyldimethylsilyl)oxy)-3a-((methyl(phenyl)amino)methyl)-3-phenyl-1,3a,4,5,6,6a-hexahydropentalen-2-yl trifluoromethanesulfonate (10):** A flame-dried round-bottom flask was charged with a stirbar and NaH (60% dispersion in mineral oil, 1.08 mmol, 44.8 mg) then evacuated and backfilled with nitrogen four times. Dry DMF (4 ml) was then added, and the reaction flask was cooled to 0 ºC. **S6** (approximately 0.52 mmol) was added slowly as a solution in dry benzene (3.4 ml) via syringe. After stirring at 0 ºC for 4 h, $PhNTf_2$ (0.81 mmol, 289.7 mg) was added as a solid and reaction put back under nitrogen. The resulting mixture was allowed to warm to 23 ºC and stirred for 16 h. The mixture was then quenched with EtOAc before exposing to atmosphere and further diluting with EtOAc and $H_2O$. The organic layer was washed four times with $H_2O$ then brine, dried over $Na_2SO_4$, and filtered. Filtrate was concentrated under reduced pressure to obtain a black oil. The crude product was purified by flash chromatography on silica with 1-10% EtOAc/hexane eluent to obtain the title compound as a yellow oil (220.8 mg, 73% over 2 steps from photoredox conjugate addition).

**$^{1}$H NMR** (600 MHz, $CDCl_3$) δ 7.44 – 7.32 (m, 5H), 7.18 (t, J = 7.7 Hz, 2H), 6.76 – 6.64 (m, 3H), 3.98 (q, J = 4.1 Hz, 1H), 3.59 (d, J = 15.3 Hz, 1H), 3.52 (d, J = 15.3 Hz, 1H), 3.02 (dd, J = 17.0, 9.9 Hz, 1H), 2.84 (s, 3H), 2.62 (dt, J = 9.8, 2.9 Hz, 1H), 2.43 (dd, J = 17.2, 2.6 Hz, 1H), 2.04 – 1.95 (m, 1H), 1.90 – 1.84 (m, 1H), 1.84 – 1.77 (m, 1H), 1.74 – 1.67 (m, 1H), 0.88 (s, 9H), 0.05 (s, 3H), 0.04 (s, 3H).

**$^{13}$C NMR** (151 MHz, $CDCl_3$) δ 150.9, 142.6, 135.3, 131.9, 129.2, 129.0, 128.7, 128.6, 118.34 (q, *J* = 320.2 Hz), 116.8, 112.8, 81.4, 61.8, 58.9, 50.4, 40.1, 35.7, 34.3, 33.0, 26.0, 18.2, -4.49, -4.51.

HPLC method D **LRMS** (ESI, APCI) m/z: calc'd for $C_{29}H_{39}F_3NO_4SSi$ $(M+H)^+$ 582.2, found 581.7.

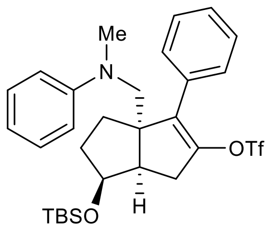

**(3a,6,6a)-6-((tert-butyldimethylsilyl)oxy)-3a-((methyl(phenyl)amino)methyl)-3-phenyl-1,3a,4,5,6,6a-hexahydropentalen-2-yl trifluoromethanesulfonate (S8):** A flame-dried round-bottom flask was charged with a stirbar and NaH (60% dispersion in mineral oil, 4.0 mmol, 160.0 mg) then evacuated and backfilled with nitrogen four times. Dry DMF (15 ml) was then added, and the reaction flask was cooled to 0 ºC. **S7** (approximately 2.0 mmol) was added slowly as a solution in dry benzene (20 ml) via syringe. After stirring at 0 ºC for 1.25 h, $PhNTf_2$ (3.0 mmol, 1.07 g) was added as a solid and reaction put back under nitrogen. The resulting mixture was allowed to warm to 23 ºC and stirred for 3 h. The mixture was then quenched with EtOAc before exposing to atmosphere and further diluting with EtOAc and $H_2O$. The organic layer was washed four times with $H_2O$ then brine, dried over $Na_2SO_4$, and filtered. Filtrate was concentrated under reduced pressure to obtain a black oil. The crude product was purified by flash chromatography on silica with 1-10% EtOAc/hexane eluent to obtain the title compound as a yellow oil (0.93 g, 80% over 2 steps from photoredox conjugate addition).

**$^{1}$H NMR** (500 MHz, $CDCl_3$) δ 7.44 – 7.31 (m, 5H), 7.20 (d, J = 7.9 Hz, 2H), 6.74 – 6.63 (m, 3H), 4.18 (dd, J = 1117.8, 6.1 Hz, 1H), 3.54 (d, J = 15.4 Hz, 1H), 3.48 (d, J = 15.3 Hz, 1H), 2.94 (d, J = 13.9 Hz, 1H), 2.92 (s, 3H), 2.74 – 2.60 (m, 2H), 2.01 – 1.90 (m, 1H), 1.83 – 1.39 (m, 3H), 0.87 (s, 9H), 0.03 (s, 3H), 0.02 (s, 3H).

**$^{13}$C NMR** (126 MHz, $CDCl_3$) δ 150.8, 144.1, 134.6, 132.0, 129.3, 129.0, 128.7, 128.4, 118.4 (q, $J$ = 320.3 Hz), 116.7, 112.3, 74.3, 61.5, 58.5, 46.1, 40.3, 34.2, 32.4, 30.2, 25.9, 18.2, -4.5, -4.9.

HPLC method D **LRMS** (ESI, APCI) m/z: calc'd for $C_{29}H_{39}F_3NO_4SSi$ $(M+H)^+$ 582.2, found 581.7.

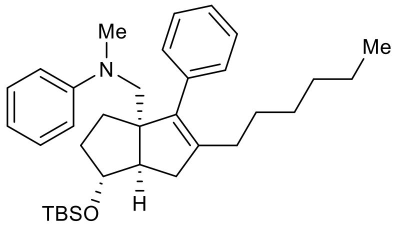

**N-(((1R,3aS,6aR)-1-((tert-butyldimethylsilyl)oxy)-5-hexyl-4-phenyl-2,3,6,6a-tetrahydropentalen-3a(1H)-yl)methyl)-N-methylaniline (14):** A round-bottom flask was charged with a stirbar and LiCl (10 mmol, 424.0 mg) then heated to 160 ºC under vacuum for 20 minutes before cooling again to 23 ºC. Once cooled, zinc (30 mesh, 15 mmol, 980.0 mg) was added and re-heated to 160 ºC under vacuum for 20 minutes. While cooling back to 23 ºC, the flask was backfilled with nitrogen and evacuated three times. Once the flask cooled, dry THF was added (10 ml) and began stirring vigorously. To the vigorously stirred suspension was added 1,2-dibromoethane (0.5 mmol, 40 µl), trimethylsilyl chloride (0.1 mmol, 13 µl), and one drop of a 1M solution of $I_2$ in dry THF under nitrogen. Once the brown color of the $I_2$ had disappeared (about 2 minutes), 1-bromohexane (10 mmol, 1.65 g) was added neat via syringe and the solution was heated to 50 ºC. After stirring at 50 ºC 16 h, a titer for the hexylzinc bromide of 0.1 M was obtained by colorimetric titration of an aliquot with a 1M solution of $I_2$ in dry THF (equivalence point reached when $I_2$ color persists with stirring). A separate flame-dried reaction vial was charged with a stirbar, **8** (0.034 mmol, 20.0 mg), $Pd(OAc)_2$ (0.44 µmol, 0.1 mg), and SPhos (1.2 µmol, 0.5 mg). The reaction vial was evacuated and backfilled with nitrogen four times then hexylzinc bromide solution added (0.041 mmol, 410 µl) via syringe. The resulting mixture was heated to 50 ºC for 16 h before cooling back to 23 ºC and pushing through a plug of silica with ethyl acetate. Filtrate concentrated under reduced pressure to a black oil. The crude mixture was taken to the next step without further purification.

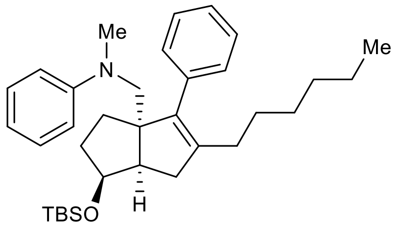

**N-(((1S,3aS,6aR)-1-((tert-butyldimethylsilyl)oxy)-5-hexyl-4-phenyl-2,3,6,6a-tetrahydropentalen-3a(1H)-yl)methyl)-N-methylaniline (S9):** A round-bottom flask was charged with a stirbar and LiCl (20 mmol, 848.0 mg) then heated to 140 ºC under vacuum for 10 minutes before cooling again to 23 ºC. Once cooled, zinc (30 mesh, 30 mmol, 1.961 g) was added and re-heated to 140 ºC under vacuum for 10 minutes. While cooling back to 23 ºC, the flask was backfilled with nitrogen and evacuated three times. Once the flask cooled, dry THF was added (20 ml) and began stirring vigorously. To the vigorously stirred suspension was added 1,2-dibromoethane (1.0 mmol, 86 µl), trimethylsilyl chloride (0.2 mmol, 25 µl), and two drops of a 1M solution of $I_2$ in dry THF under nitrogen. Once the brown color of the $I_2$ had disappeared (about 10 minutes), 1-iodohexane (20 mmol, 2.95 ml) was added neat via syringe and the solution was heated to reflux for 10 seconds then to 50 ºC. After stirring at 50 ºC for 4 h, a titer for the hexylzinc iodide of 0.67 M was obtained by colorimetric titration of an aliquot with a 1M solution of $I_2$ in dry THF (equivalence point reached when $I_2$ color persists with stirring). A separate flame-dried reaction vial was charged with a stirbar, **S8** (0.47 mmol, 272.0 mg), $Pd(OAc)_2$ (0.023 mmol, 5.2 mg), and SPhos (0.047 mol, 19.3 mg). The reaction vial was evacuated and backfilled with nitrogen four times then dry THF (0.5 ml) added and the resulting solution stirred at 23 ºC. After 10 minutes hexylzinc iodide solution added (1.88 mmol, 2.8 ml) via syringe. The resulting mixture was heated to 50 ºC for 16 h before cooling back to 23 ºC and pushing through a plug of silica with DCM. Filtrate concentrated under reduced pressure to a black oil. The crude product was taken to the next step without further purification.

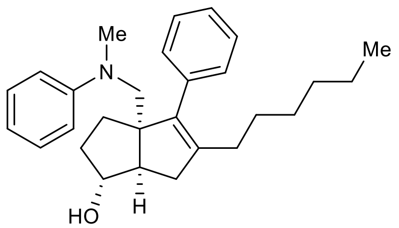

**(1,3a,6a)-5-hexyl-3a-((methyl(phenyl)amino)methyl)-4-phenyl-1,2,3,3a,6,6a-hexahydropentalen-1-ol (9):** A round-bottom flask was charged with a stirbar, **14 (**18 µmol, 9.4 mg), and 1:1 DCM:MeOH (1 ml). The resulting solution was stirred at 23 ºC and two drops of concentrated hydrochloric acid added. After 2 h the reaction was diluted with EtOAc and washed twice with saturated aqueous $NaHCO_3$, then $H_2O$, and brine. The organic layer was dried over $Na_2SO_4$, filtered and concentrated under reduced pressure. The resulting oil was put through a plug of silica with 20% EtOAc/hexane to collect the title compound (6.7 mg, 49% over 2 steps).

**$^1$H NMR** (600 MHz, $CDCl_3$) δ 7.35 (t, *J* = 7.4 Hz, 2H), 7.30 (t, *J* = 7.1 Hz, 1H), 7.17 (t, *J* = 7.7 Hz, 2H), 7.09 (d, *J* = 7.4 Hz, 2H), 6.71 – 6.64 (m, 3H), 3.94 (s, 1H), 3.48 (d, *J* = 15.2 Hz, 1H), 3.40 (d, *J* = 15.2 Hz, 1H), 3.00 (s, 3H), 2.79 (dd, *J* = 17.1, 9.5 Hz, 1H), 2.56 (d, *J* = 9.5 Hz, 1H), 2.14 (dd, *J* = 17.2, 2.8 Hz, 1H), 1.90 (t, *J* = 7.6 Hz, 2H), 1.90 – 1.64 (m, 4H), 1.37 – 1.29 (m, 2H), 1.29 – 1.20 (m, 2H), 1.20 – 1.12 (m, 4H), 0.85 (t, *J* = 7.2 Hz, 3H).

**$^{13}$C NMR** (151 MHz, $CDCl_3$) δ 151.4, 140.6, 140.2, 138.0, 130.0, 129.2, 128.2, 126.8, 116.9, 113.0, 81.5, 66.6, 59.1, 53.1, 41.8, 39.4, 34.0, 31.8, 30.9, 29.3, 29.2, 28.1, 22.7, 14.2.

HPLC method F **LRMS** (ESI, APCI) m/z: calc'd for $C_{28}H_{38}NO$ $(M+H)^+$ 404.3, found 403.9.

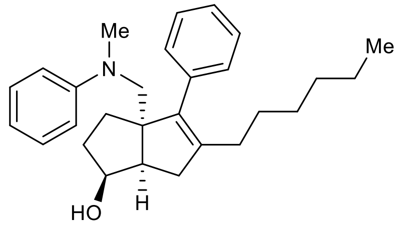

**(1,3a,6a)-5-hexyl-3a-((methyl(phenyl)amino)methyl)-4-phenyl-1,2,3,3a,6,6a-hexahydropentalen-1-ol (S10):** A round-bottom flask was charged with a stirbar, **S11** (approx. 0.47 mmol), and 1:1 DCM:MeOH (15 ml). The resulting solution was stirred at 23 ºC and four drops of concentrated hydrochloric acid added. After 1 h the reaction was diluted with EtOAc and washed three times with $H_2O$ then brine. The organic layer was dried over $Na_2SO_4$, filtered and concentrated under reduced pressure to collect crude. The crude material was then loaded on silica and eluted with 5-15% EtOAc/hexanes to collect the title compound (82.0 mg, 43% over 2 steps.).

**$^1$H NMR** (600 MHz, $CDCl_3$) δ 7.35 (t, J = 7.5 Hz, 2H), 7.29 (t, J = 7.4 Hz, 1H), 7.15 (t, J = 7.1 Hz, 2H), 7.12 (d, J = 7.2 Hz, 2H), 6.64 (t, J = 7.2 Hz, 1H), 6.59 (d, J = 8.3 Hz, 2H), 4.19 (p, J = 6.3 Hz, 1H), 3.46 (d, J = 15.3 Hz, 1H), 3.39 (d, J = 15.3 Hz, 1H), 2.96 (s, 3H), 2.73 (t, J = 6.8 Hz, 1H), 2.65 (d, J = 17.1 Hz, 1H), 2.47 (dd, J = 17.2, 9.2 Hz, 1H), 2.02 – 1.91 (m, 2H), 1.90 – 1.82 (m, 1H), 1.76 (dt, J = 12.4, 5.2 Hz, 1H), 1.66 – 1.55 (m, 2H), 1.42 – 1.36 (m, 3H), 1.28 – 1.22 (m, 2H), 1.24 – 1.14 (m, 3H), 0.85 (t, J = 7.2 Hz, 3H).

**$^{13}$C NMR** (151 MHz, $CDCl_3$) δ 151.0, 142.4, 140.6, 137.8, 130.0, 129.1, 128.2, 126.8, 116.0, 111.9, 75.2, 66.5, 58.6, 47.2, 40.6, 34.1, 33.2, 31.8, 30.8, 29.5, 29.3, 28.2, 22.8, 14.2.

HPLC method E **LRMS** (ESI, APCI) m/z: calc'd for $C_{28}H_{38}NO$ $(M+H)^+$ 404.3, found 403.9.

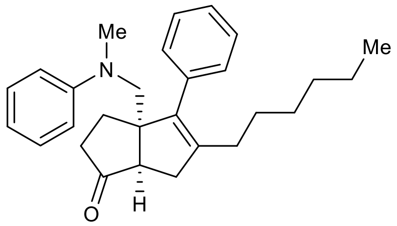

**(3a,6a)-5-hexyl-3a-((methyl(phenyl)amino)methyl)-4-phenyl-3,3a,6,6a-tetrahydropentalen-1(2H)-one (S11):** A 1-dram reaction vial was charged with a stirbar, **S10** (0.32 mmol, 131.0 mg), and MeCN (3 ml). The resulting solution stirred at 23 ºC then TPAP (0.032 mmol, 11.4 mg) and NMO (3.24 mmol, 379.6 mg) added. The reaction solution continued to stir for 40 minutes before eluting through a plug of silica. The resulting crude material was then loaded on silica and eluted with 10% EtOAc/hexanes to collect the title compound (109.6 mg, 84%).

**$^1$H NMR** (600 MHz, $CDCl_3$) δ 7.39 (t, *J* = 6.9 Hz, 2H), 7.34 (t, *J* = 7.1 Hz, 1H), 7.18 (t, *J* = 7.9 Hz, 2H), 7.04 (d, *J* = 6.7 Hz, 2H), 6.68 (t, *J* = 7.1 Hz, 1H), 6.64 (d, *J* = 8.3 Hz, 2H), 3.63 (d, *J* = 15.5 Hz, 1H), 3.45 (d, *J* = 15.5 Hz, 1H), 3.00 (s, 2H), 2.78 (dd, *J* = 16.4, 7.6 Hz, 1H), 2.64 (d, *J* = 7.6 Hz, 1H), 2.61 (d, *J* = 16.5 Hz, 1H), 2.34 (dd, *J* = 18.5, 8.8 Hz, 1H), 2.24 – 2.12 (m, 1H), 2.07 – 1.88 (m, 4H), 1.33 (p, *J* = 7.2 Hz, 2H), 1.29 – 1.19 (m, 2H), 1.19 – 1.10 (m, 4H), 0.84 (t, *J* = 7.2 Hz, 3H).

**$^{13}$C NMR** (126 MHz, $CDCl_3$) δ 222.8, 150.4, 143.8, 139.7, 136.7, 129.4, 129.2, 128.5, 127.2, 116.5, 111.8, 64.3, 57.8, 53.2, 41.6, 37.5, 36.8, 31.6, 29.4, 29.0, 27.8, 26.8, 22.7, 14.1.

HPLC method D **LRMS** (ESI, APCI) m/z: calc'd for $C_{28}H_{36}NO$ $(M+H)^+$ 402.3, found 401.9.

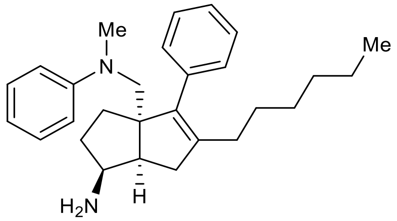

**(1,3a,6a)-5-hexyl-3a-((methyl(phenyl)amino)methyl)-4-phenyl-1,2,3,3a,6,6a-hexahydropentalen-1-amine (S12):** A 1-dram reaction vial was charged with a stirbar, **S11** (0.25 mmol, 98.2 mg), and EtOH (2.5 ml). A solution of $NH_3$ in MeOH (7N, 1.22 mmol, 175 µl) followed immediately by $Ti(O^iPr)_4$ (0.37 mmol, 111 µl) was added and the vial sealed. The resulting solution was stirred at 23 ºC for 6 h before unsealing vial and adding $NaBH_4$ (0.73 mmol, 27.6 mg) and continuing stirring at 23 ºC for 16 h. Reaction was then diluted with EtOAc and saturated aqueous Rochelle's salt and sonicated for 5 min. The resulting slurry was washed twice with saturated aqueous Rochelle's salt, twice with $H_2O$, then brine. The organic layer was dried over $Na_2SO_4$, filtered, and filtrate concentrated under reduced pressure to collect the crude material (7.5:1 dr *endo*:*exo*). Crude material purified by flash chromatography on silica with

20:80:0 to 99:0:1 EtOAc:hexanes:$Et_3N$ eluent to collect the title compound as a 7.5:1 *endo*:*exo* mix of diastereomers (33.5 mg, 34% combined diastereomers).

*endo* diastereomer: $^1$**H NMR** (600 MHz, $CDCl_3$) δ 7.36 – 7.32 (m, 2H), 7.31 – 7.27 (m, 1H), 7.16 – 7.12 (m, 2H), 7.11 – 7.08 (m, 2H), 6.63 (t, J = 7.2 Hz, 1H), 6.57 (d, J = 8.3 Hz, 2H), 3.47 (d, J = 15.4 Hz, 1H), 3.37 (d, J = 15.3 Hz, 1H), 3.30 (q, J = 8.5 Hz, 1H), 2.95 (s, 3H), 2.68 (td, J = 8.3, 4.0 Hz, 1H), 2.51 – 2.47 (m, 2H), 1.98 – 1.92 (m, 2H), 1.87 – 1.80 (m, 1H), 1.72 – 1.67 (m, 1H), 1.55 (td, J = 12.2, 6.0 Hz, 1H), 1.41 – 1.33 (m, 3H), 1.28 – 1.22 (m, 3H), 1.21 – 1.15 (m, 3H), 0.85 (t, J = 7.2 Hz, 3H).

HPLC method F **LRMS** (ESI, APCI) m/z: calc'd for $C_{28}H_{39}N_2$ $(M+H)^+$ 403.3, found 403.0.

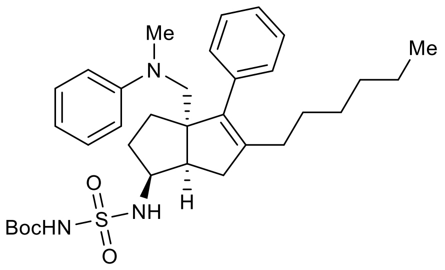

**tert-butyl (N-((1,3a,6a)-5-hexyl-3a-((methyl(phenyl)amino)methyl)-4-phenyl-1,2,3,3a,6,6a-hexahydropentalen-1-yl)sulfamoyl)carbamate (S13):** An oven-dried vial was charged with a stirbar, $^t$BuOH (1.44 mmol, 106.7 mg), and DCM (1.57 ml) then evacuated under reduced pressure and backfilled with nitrogen three times and cooled to 0 ºC. Chlorosulfonyl isocyanate (1.3 mmol, 113 µl) was then added dropwise via syringe and the solution allowed to warm to 23 ºC over 35 minutes. A 100 µl portion of this solution was added slowly via syringe to a solution of **S12** (0.083 mmol, 33.5 mg. 7.5:1 *endo*:*exo*) and $Et_3N$ (0.125 mmol, 17 µl) in DCM (100 µl) at 0 ºC under nitrogen. This combined solution was allowed to warm to 23 ºC gradually over 3.75 h then diluted with EtOAc. The diluted solution was washed three times with 0.5 M HCl then $H_2O$ and brine. The organic layer was dried over $Na_2SO_4$, filtered, and filtrate concentrated under reduced pressure to collect the crude material. Crude material purified by flash chromatography on silica with 10-50% EtOAc/hexanes to collect impure product taken to the next step without further purification.

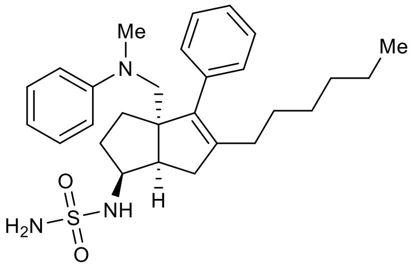

**N-((1,3a,6a)-5-hexyl-3a-((methyl(phenyl)amino)methyl)-4-phenyl-1,2,3,3a,6,6a-hexahydropentalen-1-yl)sulfamide (15):** A solution of 3:1 dioxane/concentrated aqueous HCl was frozen in an ice bath then allowed to slowly warm to 23 ºC. As soon as the entire solution had re-melted, 0.8 ml was transferred to a chilled (~0 ºC, but NOT in an ice bath) vial containing a stirbar and **S13** (38 µmol, 22.2 mg). The solution was allowed to slowly warm to 23 ºC and continue reacting for 20 h until **S13** was consumed. The reaction solution was diluted with EtOAc and washed four times with $H_2O$ then twice with brine. The organic layer was dried over $Na_2SO_4$, filtered, and filtrate concentrated under reduced pressure to collect the crude material. This crude material was purified by flash chromatography on silica with 5-30% EtOAc/hexanes then by preparative HPLC with 40 ml/min 50-70% MeCN/$H_2O$ gradient over 25 minutes to collect the title compound as a single diastereomer (10.4 mg, 26% over 2 steps).

**$^1$H NMR** (600 MHz, $CDCl_3$) δ 7.36 (t, J = 6.9 Hz, 2H), 7.31 (t, J = 7.7 Hz, 1H), 7.15 (t, J = 6.9 Hz, 2H), 7.09 (d, J = 6.9 Hz, 2H), 6.65 (t, J = 7.2 Hz, 1H), 6.59 (d, J = 8.3 Hz, 2H), 4.38 (s, 2H), 4.32 (d, J = 8.0 Hz, 1H), 3.85 – 3.75 (m, 1H), 3.51 (d, J = 15.4 Hz, 1H), 3.38 (d, J = 15.4 Hz, 1H), 2.96 (s, 3H), 2.93 (td, J = 8.7, 3.1 Hz, 1H), 2.59 (dd, J = 17.4, 9.0 Hz, 1H), 2.44 (dd, J = 17.4, 3.1 Hz, 1H), 2.02 – 1.92 (m, 3H), 1.79 – 1.73 (m, 1H), 1.66 – 1.56 (m, 2H), 1.38 (p, J = 7.1 Hz, 2H), 1.28 – 1.22 (m, 2H), 1.21 – 1.14 (m, 4H), 0.85 (t, J = 7.2 Hz, 3H).

**$^{13}$C NMR** (151 MHz, $CDCl_3$) δ 150.9, 141.9, 140.6, 137.3, 129.8, 129.2, 128.3, 127.0, 116.3, 112.1, 66.4, 58.4, 57.5, 46.1, 40.9, 34.8, 32.3, 31.7, 31.1, 29.4, 29.3, 28.2, 22.7, 14.2.

HPLC method G **LRMS** (ESI, APCI) m/z: calc'd for $C_{28}H_{39}N_3O_2S$ $(M+H)^+$ 482.3, found 481.9.

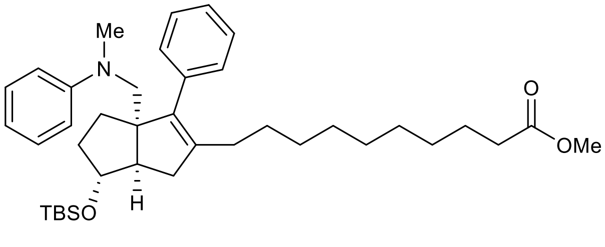

**methyl 10-((3a,6,6a)-6-((tert-butyldimethylsilyl)oxy)-3a-((methyl(phenyl)amino)methyl)-3-phenyl-1,3a,4,5,6,6a-hexahydropentalen-2-yl)decanoate (11):** A round-bottom flask was charged with a stirbar and LiCl (8 mmol, 339.0 mg) then heated to 140 ºC under vacuum for 20 minutes before cooling again to 23 ºC. Once cooled, zinc (30 mesh, 12 mmol, 784.6 mg) was added and re-heated to 140 ºC under vacuum for 1 h. While cooling back to 23 ºC, the flask was backfilled with nitrogen and evacuated three times. Once the flask cooled, dry THF was added (8 ml) and began stirring vigorously. To the vigorously stirred suspension was added 1,2-dibromoethane (0.4 mmol, 35 µl), trimethylsilyl chloride (0.08 mmol, 10 µl), and one drop of a 1M solution of $I_2$ in dry THF under nitrogen. Once the brown color of the $I_2$ had disappeared (about 2.5 h), methyl 10-iodododecanoate (8 mmol, 2.5 g) was added neat via syringe and the solution was heated to 50 ºC. After stirring at 50 ºC 16 h, a titer for the alkylzinc iodide of 0.78 M was obtained by colorimetric titration of an aliquot with a 1M solution of $I_2$ in dry THF (equivalence point reached when $I_2$ color persists with stirring). A separate flame-dried reaction vial was charged with a stirbar, **10** (0.34 mmol, 200.0 mg), Sphos G3 (17 µmol, 13.3 mg), and SPhos (34 µmol, 13.9 mg). The reaction vial was evacuated and backfilled with nitrogen four times then THF (0.5 ml) added and began heating to 50 ºC. After 10 minutes, alkylzinc iodide solution added (1.7 mmol, 2.1 ml) via syringe. The resulting mixture continued to stir at 50 ºC for 40 h before cooling back to 23 ºC and pushing through a plug of silica with ethyl acetate. Filtrate concentrated under reduced pressure to a black oil. The crude product was taken to the next step without further purification.

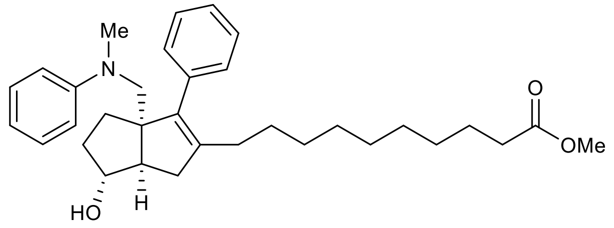

**methyl 10-((3a,6,6a)-6-hydroxy-3a-((methyl(phenyl)amino)methyl)-3-phenyl-1,3a,4,5,6,6a-hexahydropentalen-2-yl)decanoate (12):** A round-bottom flask was charged with a stirbar, **11** (approx. 0.34 mmol), and MeOH (12 ml). The resulting solution was stirred at 23 ºC and two drops of concentrated hydrochloric acid added. After 16 h the reaction was diluted with EtOAc and washed with saturated aqueous $NaHCO_3$, $H_2O$ twice, then brine. The organic layer was dried over $Na_2SO_4$, filtered and concentrated under reduced pressure to collect a crude mixture. The crude mixture was purified by flash chromatography over silica with 10-50% EtOAc/hexanes eluent to collect the title compound (94.0 mg, 54% over two steps).

**$^1$H NMR** (600 MHz, $CDCl_3$) δ 7.35 (t, J = 7.4 Hz, 2H), 7.32 – 7.27 (m, 1H), 7.17 (t, J = 7.4 Hz, 2H), 7.09 (d, J = 7.1 Hz, 2H), 6.71 – 6.65 (m, 3H), 3.95 – 3.91 (m, 1H), 3.66 (d, J = 0.7 Hz, 3H), 3.48 (d, J = 15.2 Hz, 1H), 3.39 (d, J = 15.2 Hz, 1H), 3.00 (s, 3H), 2.78 (dd, J = 17.1, 9.5 Hz, 1H), 2.56 (d, J = 9.4 Hz, 1H), 2.29 (t, J = 7.5 Hz, 2H), 2.14 (dd, J = 17.2, 2.9 Hz, 1H), 1.90 (t, J = 7.6 Hz, 2H), 1.88 – 1.82 (m, 1H), 1.79 – 1.64 (m, 3H), 1.60 (p, J = 7.4 Hz, 2H), 1.36 – 1.12 (m, 12H).

**$^{13}$C NMR** (126 MHz, $CDCl_3$) δ 174.5, 151.3, 140.5, 140.3, 138.0, 129.9, 129.1, 128.8, 126.8, 116.8, 112.9, 81.4, 66.6, 59.1, 53.1, 51.6, 41.8, 39.3, 34.2, 34.0, 30.9, 29.46, 29.43, 29.42, 29.35, 29.25, 29.23, 28.0, 25.1.

HPLC method F **LRMS** (ESI, APCI) m/z: calc'd for $C_{33}H_{46}NO_3$ $(M+H)^+$ 504.4, found 503.9.

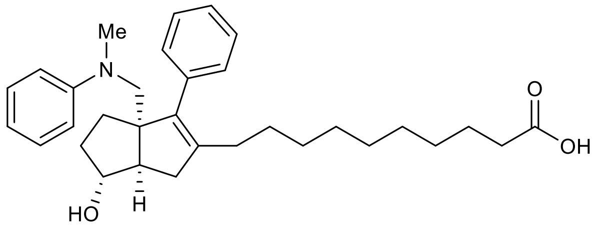

**10-((3a,6,6a)-6-hydroxy-3a-((methyl(phenyl)amino)methyl)-3-phenyl-1,3a,4,5,6,6a-hexahydropentalen-2-yl)decanoic acid (13):** A 1-dram vial was charged with a stirbar, **12** (0.03 mmol, 15.0 mg), $LiOH{\cdot}H_2O$ (0.3 mmol, 12.5 mg), and 0.3 ml of 5:1 THF/$H_2O$ solution. The resulting suspension was stirred at 50 ºC for 16 h. The reaction was then diluted with EtOAc and $H_2O$. The aqueous layer was brought to pH ~0 and extracted three times with EtOAc. The EtOAc layers were then combined, washed twice with brine, dried over $Na_2SO_4$, filtered and concentrated under reduced pressure to afford the title compound (14.6 mg, quant.).

**$^1$H NMR** (600 MHz, $CDCl_3$) δ 7.35 (t, J = 7.4 Hz, 2H), 7.32 – 7.27 (m, 1H), 7.17 (t, J = 8.0 Hz, 2H), 7.09 (d, J = 7.3 Hz, 2H), 6.71 – 6.65 (m, 3H), 3.94 (s, 1H), 3.47 (d, J = 15.1 Hz, 1H), 3.39 (d, J = 15.3 Hz, 1H), 2.99 (s, 3H), 2.78 (dd, J = 17.1, 9.5 Hz, 1H), 2.56 (d, J = 9.5 Hz, 1H), 2.33 (t, J = 7.5 Hz, 2H), 2.14 (dd, J = 17.2, 2.9 Hz, 1H), 1.90 (t, J = 7.6 Hz, 2H), 1.88 – 1.82 (m, 1H), 1.81 – 1.65 (m, 3H), 1.62 (p, J = 7.4 Hz, 2H), 1.37 – 1.12 (m, 12H).

**$^{13}$C NMR** (151 MHz, $CDCl_3$) δ 178.9, 151.4, 140.5, 140.3, 138.0, 129.9, 129.2, 128.2, 126.8, 116.9, 113.0, 81.5, 66.6, 59.1, 53.1, 41.8, 39.3, 34.0, 30.9, 29.42, 29.40, 29.39, 29.3, 29.22, 29.16, 28.0, 24.8.

HPLC method F **LRMS** (ESI, APCI) m/z: calc'd for $C_{32}H_{44}NO_3$ $(M+H)^+$ 490.3, found 489.9.

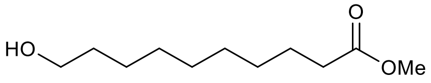

**methyl 10-hydroxydecanoate (S14):** A round-bottom flask was charged with a stirbar, 10-hydroxydecanoic acid (15.93 mmol, 3.0 g), and MeOH (300 ml). The resulting solution was charged with three drops of concentrated aqueous HCl and stirred at 50 ºC for 2 h. The reaction was then concentrated under reduced pressure and taken to the next step without further purification.

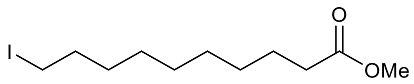

**methyl 10-iodododecanoate** (**S15**)**:** A flame-dried round-bottom flask was charged with **S14** (approximately 15.93 mmol), $PPh_3$ (31.86 mmol, 8.356 g), and imidazole (47.79 mmol, 3.253 g). The flask was then evacuated under reduced pressure and backfilled with nitrogen three times. DCM (75 ml) was then added and the solution cooled to 0 ºC. A solution of $I_2$ (31.86 mmol, 8.086 g) in THF (35 mL) was then added slowly via syringe and the resulting solution was stirred for 16 h after warming to 23 ºC. The reaction was then diluted with EtOAc and washed once with aqueous 10% $Na_2S_2O_3$, three times with $H_2O$, and twice with brine. The organic layer was dried over $Na_2SO_4$, filtered and concentrated under reduced pressure to afford a white solid. The crude solid was purified by flash chromatography over silica with 10-50% EtOAc/hexanes eluent to collect the title compound (4.904 g, 99% over two steps).

**$^1$H NMR** (600 MHz, $CDCl_3$) δ 3.67 (s, 3H), 3.18 (t, $J$ = 7.0 Hz, 2H), 2.30 (t, $J$ = 7.6 Hz, 2H), 1.81 (p, $J$ = 7.2 Hz, 2H), 1.62 (p, $J$ = 7.2 Hz, 2H), 1.38 (p, $J$ = 6.6 Hz, 2H), 1.29 (b, 8H).

(10) The PyMOL Molecular Graphics System, Version 2.0 Schrödinger, LLC.
